# Insulin/insulin-like growth factor signaling pathway promotes higher fat storage in *Drosophila* females

**DOI:** 10.1101/2024.11.18.623936

**Authors:** Puja Biswas, Jennifer A. Bako, J. Beatrice Liston, Huaxu Yu, Lianna W. Wat, Colin J. Miller, Michael D. Gordon, Tao Huan, Molly Stanley, Elizabeth J. Rideout

## Abstract

In *Drosophila*, adult females store more fat than males. While the mechanisms that restrict body fat in males are becoming clearer, less is known about how females achieve higher fat storage. Here, we perform a detailed investigation of the mechanisms that promote higher fat storage in females. We show greater intake of dietary sugar supports higher fat storage due to female-biased remodeling of the fat body lipidome. Dietary sugar stimulates a female-specific increase in *Drosophila* insulin-like peptide 3 (Dilp3), which acts together with greater peripheral insulin sensitivity to augment insulin/insulin-like growth factor signaling pathway (IIS) activity in adult females. Indeed, Dilp3 overexpression prevented the female-biased decrease in body fat after removal of dietary sugar. Given that adult-specific IIS inhibition caused a female-biased decrease in body fat, our data reveal IIS as a key determinant of female fat storage.

## Introduction

In *Drosophila*, as in other animals, triglyceride is the main stored form of fat^1^. The primary depot for stored fat in insects is called the fat body^1^; however, other cell types also synthesize and store triglyceride (*e.g.*, gut, oenocytes, neurons, glia, muscle, testis, ovary)^2–9^. While many factors influence the total amount of stored fat, one important determinant of body fat in flies is biological sex^5,9,10^. For example, *Drosophila* virgin females store more fat than males^5^, a difference that is amplified post-mating^9,11^. Virgin females also break down fat more slowly than males after nutrient withdrawal^5^. These sex differences in body fat regulation are physiologically significant: females use their greater fat stores to support the energetic demands of reproduction^9,12^ and to survive long bouts of nutrient deprivation^5^, whereas males need to restrict excess triglyceride accumulation to maintain fertility^8^.

Over the past two decades, studies in *Drosophila* have played a key role in identifying many genes and pathways that contribute to whole-body fat storage. For example, loss-of-function experiments with genes predicted to promote triglyceride synthesis lead to reduced fat storage in both individual cells and in the whole body^12–16^. In contrast, genetically inhibiting genes that facilitate triglyceride breakdown significantly augment fat storage at the cellular and organismal levels^17–19^. Gain- and loss-of-function experiments with genes predicted to regulate lipid droplet dynamics, a specialized organelle dedicated to triglyceride storage, similarly influence cellular and organismal fat stores^16,19–24^. Beyond these enzymes directly involved in triglyceride metabolism and lipid droplet biology, large-scale screening efforts have revealed many additional genes that influence *Drosophila* body fat^4,18,25–39^.

In addition to the action of individual genes, a large body of literature has identified hormones and peptides that influence whole-body fat storage^9,40–57^. Often, these hormones modulate whole-body fat storage via effects on cells that produce adipokinetic hormone (Akh)^43–46,58^ and *Drosophila* insulin-like peptides (Dilps)^30,43,45,59–62^, where Akh and most Dilps have opposing effects on whole-body fat storage^63^. For example, when nutrients are plentiful, Dilps 2, 3, and 5 are released from the insulin-producing cells (IPC) in the *Drosophila* brain into the hemolymph^43,59^. Binding of the Dilps to the insulin receptor (InR) on the fat body stimulates the activity of the insulin/insulin-like growth factor signaling pathway (IIS) to promote fat storage^53^. When nutrients are scarce, Akh is released into the hemolymph from the Akh-producing cells (APC) in the *corpora cardiaca*^46,64–67^. Akh binds to the Akh receptor located in the fat body where it promotes fat breakdown^55–57^. Despite this advanced knowledge of factors that regulate fat storage, most studies used single- or mixed-sex animal groups. As a result, less is known about the mechanisms underlying sex differences in fat storage.

Recent studies have begun to close this knowledge gap. Both large-scale RNAseq data and targeted gene expression studies reveal profound sex differences in expression of genes that regulate cellular and whole-body fat storage^5,68,69^. For example, male flies have higher mRNA levels of the *Drosophila* homolog of adipose triglyceride lipase called *brummer* (*bmm*), an enzyme that limits triglyceride accumulation across most cell types^17^. This male-biased *bmm* expression is physiologically significant, as whole-body loss of *bmm* abolishes the sex difference in fat storage via a male-biased increase in body fat^5^. A similar male bias has also been reported in the production and secretion of Akh^10^, which prevents fat accumulation to restrict body fat in males^10^. A male bias in two central lipolytic pathways therefore plays a key role in establishing the sex difference in body fat. Yet, the mechanisms that allow increased fat storage in females are less clear, as females are largely unaffected by loss of *bmm* and Akh^5,10^.

In mated females, higher levels of steroid hormone ecdysone enhance body fat by stimulating food intake^9^. Mated females also show higher mRNA levels of *dilp3*^70^, where *dilp3* has been implicated in promoting fat storage^43,62^. Whether these mechanisms also influence the sex difference in fat storage remains unclear. We therefore performed a detailed analysis of adult virgin males and females to explore how females achieve a higher level of fat storage. We found that virgin female flies normally consume more food than males, and show that female fat storage is sensitive to changes in food quantity. In particular, consumption of dietary sugar and not simply a greater number of calories explained this higher fat storage in females. Unbiased lipidomic and metabolomic analysis of abdominal carcasses from adult flies revealed dietary sugar caused a profoundly female-biased metabolic remodeling of energy-storing cells including the fat body. Given that ecdysone did not promote increased body fat in virgin females, we characterized sex differences in IIS, the other key pathway that promotes fat storage. We identified significant female biases in Dilp3 levels and fat body insulin sensitivity, leading to higher IIS activity in the female fat body. These sex biases were significant: Dilp3 overexpression in females bypassed the reduction in body fat caused by a low-sugar diet, and ablating the IPC in adults caused a female-specific reduction in body fat. Together, these data identify IIS as a key pathway that promotes higher fat storage in adult females.

## Results

### Dietary sugar promotes increased fat storage in adult females

To determine whether unmated females have higher food intake than males we used the capillary feeder (CAFE) assay^71^ to assess food consumption in 5-day-old *Drosophila* virgin males and females. Five-day-old adult *Canton-S* (*CS*) females consumed significantly more food per body weight over 24 hr than genotype-matched males (Figure 1A), a phenotype we reproduced in 5-day-old *w^11^*^18^ adults (Figure S1A). Considering changes in food intake affect body fat^26,47,72^, this suggests higher food consumption may explain this increased fat storage in females. To monitor body fat, we used a coupled colorimetric assay to measure whole-body triglyceride in male and female flies^4,35,73^, where triglyceride content was normalized to mass to account for sex differences in body size as described previously^5,10^. All non-normalized triglyceride data are available in Supplemental Table S1. Importantly, coupled colorimetric assays have been shown to accurately quantify stored triglyceride in *Drosophila*^74^. Reducing nutrient content in the diet significantly reduced body fat in *CS* females (Figure 1B). Because body fat in *CS* males was unaffected by an identical food dilution (Figure 1B), the sex difference in body fat was reduced (Figure 1B’). We next tested the contribution of specific diet components to higher fat storage in females. Given that dietary sugar affects body fat in a mixed-sex population of *Drosophila*^75–79^, we transferred newly-eclosed *CS* and *w^11^*^18^ flies to a diet lacking added sugar (0S) for 5 days^80^. In both *w^111^*^8^ and *CS* strains, adult females transferred to the 0S diet had significantly less body fat compared with genotype-matched flies kept on a widely-used lab diet with sugar (1S; Figure 1C and S1B)^81^. Because the magnitude of the reduction in body fat was smaller in *w^1118^* males (Figure 1C; sex:genotype interaction *p*<0.0001), with the same trend in *CS* males (Figure S1B), the sex difference in fat storage was reduced (Figure 1C’).

**Figure 1.**
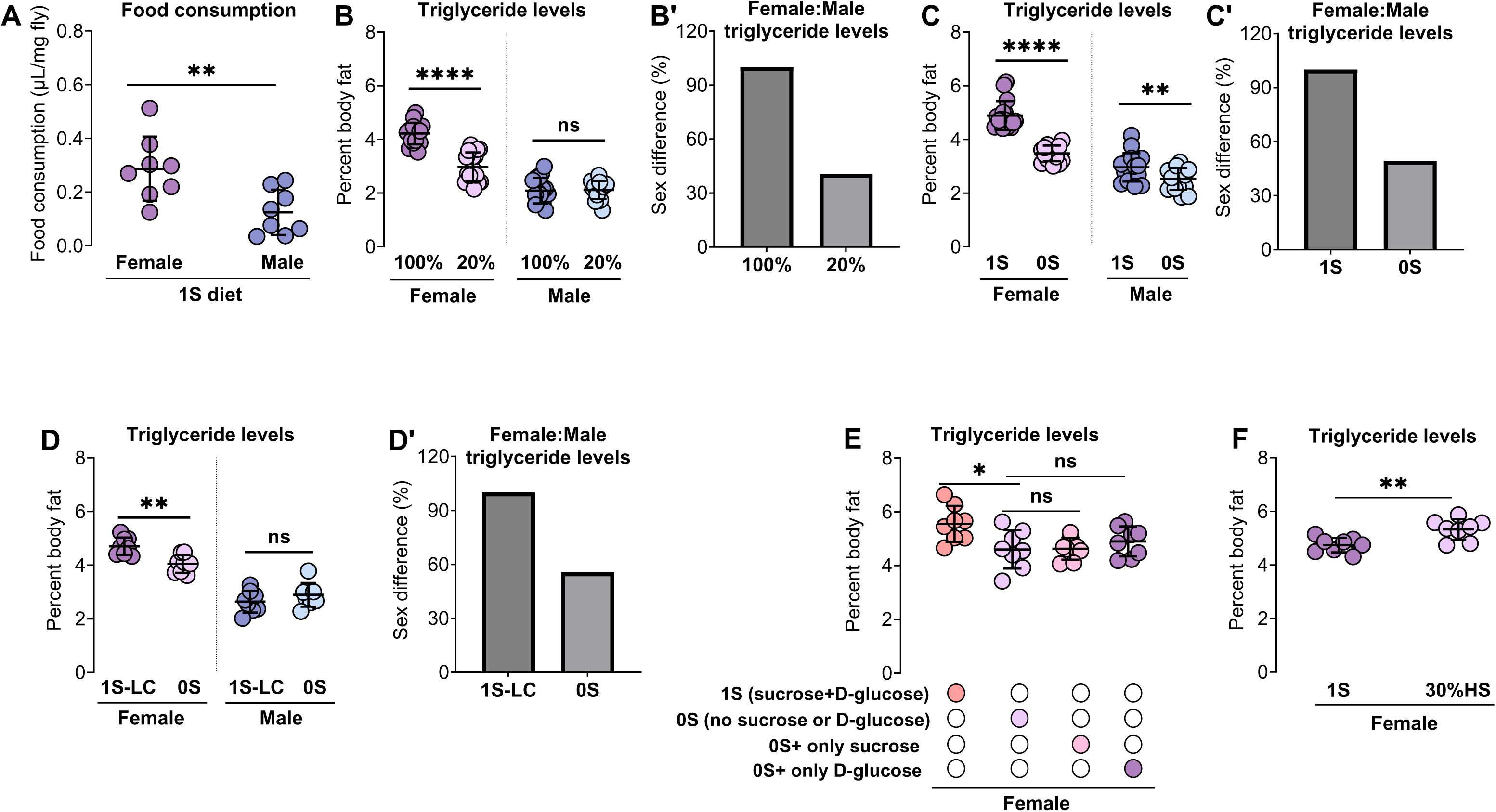
Dietary sugar promotes increased fat storage in adult females. (A) Food consumption per unit of body weight was significantly higher in 5-day-old adult *Canton-S (CS)* females than males (*p*=0.0072; Student’s *t*-test); n=8 biological replicates. (B) Whole-body triglyceride levels were significantly lower in 5-day-old *CS* females maintained on a diet with 20% of normal nutrient content compared with females kept on our normal diet (100% diet; *p<*0.0001). No change in whole-body triglyceride was observed in male flies (*p*>0.9999; sex:diet interaction *p*<0.0001). Two-way ANOVA followed by Bonferroni post-hoc test; n=16 biological replicates. (B’) Sex difference in body fat represented as a percent of the male-female difference in our normal diet. (C) Whole-body triglyceride levels were significantly higher in 5-day-old adult *w*^1118^ females and males maintained on the 1S diet compared with sex-matched flies kept on the 0S diet (Females: *p*<0.0001; Males: *p*=0.0068). The magnitude of the decrease in body fat was greater in females than in males (sex:diet interaction *p*<0.0001). Two-way ANOVA followed by Tukey HSD on data processed using the aligned rank transform for non-parametric data; n=16 biological replicates. (C’) Sex difference in body fat represented as a percent of the male-female difference in the 1S diet. (D) Whole-body triglyceride levels were significantly higher in 5-day-old adult *w*^1118^ females maintained on sugar-containing diet with reduced calories (1S-LC) compared with females kept on the 0S diet (same calorie content as 1S-LC) (*p*=0.0032); no significant change in body fat was observed in males (*p*=0.3715; sex:diet interaction *p*=0.0019). Two-way ANOVA followed by Bonferroni post-hoc test; n=8 biological replicates. (D’) Sex difference in body fat represented as a percent of the male-female difference in the lower-calorie 1S diet. (E) Whole-body triglyceride levels were not significantly different between 5-day-old adult *w^1118^* females maintained on the 0S diet and sex-matched flies kept on the 0S diet with either added 20.5 g/L of sucrose or 70.9 g/L of D-glucose diet (females: *p*=0.9998 (sucrose), *p*=0.7532 (D-glucose). One-way ANOVA followed by Tukey HSD test; n=8 biological replicates. (F) Whole-body triglyceride levels were significantly higher in 5-day-old adult *w^1118^* females maintained on our lab diet with 30% added sugar (HS) compared with females kept on the 1S diet (*p*=0.0036; Student’s *t*-test). n=8 biological replicates. All data plotted as mean ± SEM. ns indicates not significant with *p*>0.05; * *p*<0.05, ** *p*<0.01, *** *p*<0.001, **** *p*<0.0001. See also Figure S1.

The female-biased reduction in fat storage on 0S cannot be attributed to altered food intake on the low-sugar diet, as food consumption was not different between flies kept on the 0S diet compared with flies maintained on the 1S diet (Figure S1C). A decrease in calories similarly cannot explain the effect of reduced sugar on body fat, as female flies transferred to a reduced-calorie diet with sugar still had higher body fat than females kept on the 0S diet (Figure 1D and 1D’). Because adding the quantity of either sucrose (20.5 g/L) or D-glucose (70.9 g/L) that we usually put in 1S food to the 0S diet was not sufficient to restore fat storage to the same level as females maintained on the 1S diet (Figure 1E), our data suggests it is the total amount of sugar rather than a specific sugar in the diet that promotes increased fat storage in female flies. Indeed, providing females maintained on 1S with additional sugar was sufficient to further increase body fat (Figure 1F). Because the magnitude of these sugar-dependent changes in body fat (−1% or +1% body fat) are similar to the change in body fat we observe in females reared on a high-fat diet (+1%)^5^, where a high-fat diet is associated with a number of adverse physiological effects^82–84^ and changes in fertility and lifespan^85,86^, the change in body fat we observed on the 0S diet is physiologically relevant. Finally, supporting a model in which the nutritive value of sugar rather than sweet taste promotes increased female fat storage, body fat was not higher in either male or female flies kept on the 0S diet containing a non-nutritive but sweet-tasting sugar^87^ (Figure S1D). Thus, our data suggest that higher food intake, specifically increased consumption of dietary sugar, promotes higher fat storage in females.

### Female-biased effect of dietary sugar on metabolism

To assess how sex and dietary sugar affect the lipidome and metabolome of fat-storing organs in *Drosophila*, we performed untargeted mass spectrometry (MS)-based lipidomic profiling on abdominal carcasses isolated from adult male and female flies transferred to either a 1S or 0S diet as adults. We detected 464 lipid species in total (Supplemental Table S1). We first compared lipid abundance between males and females. We identified sex differences in the abundance of 163 lipids in flies maintained on the 1S diet (*p*<0.01): 117 lipid species were significantly higher in females and 46 lipid species were significantly higher in males (Figure 2A). When we examined the sex bias of lipids within each lipid subclass, we found that triglyceride was the subclass with the most species showing a female bias. Indeed, 48/57 (86%) of sex-biased triglyceride species showed a female bias in abundance (Figure 2B). This validates data from our coupled colorimetric assays, strongly supporting increased stored fat in females.

**Figure 2.**
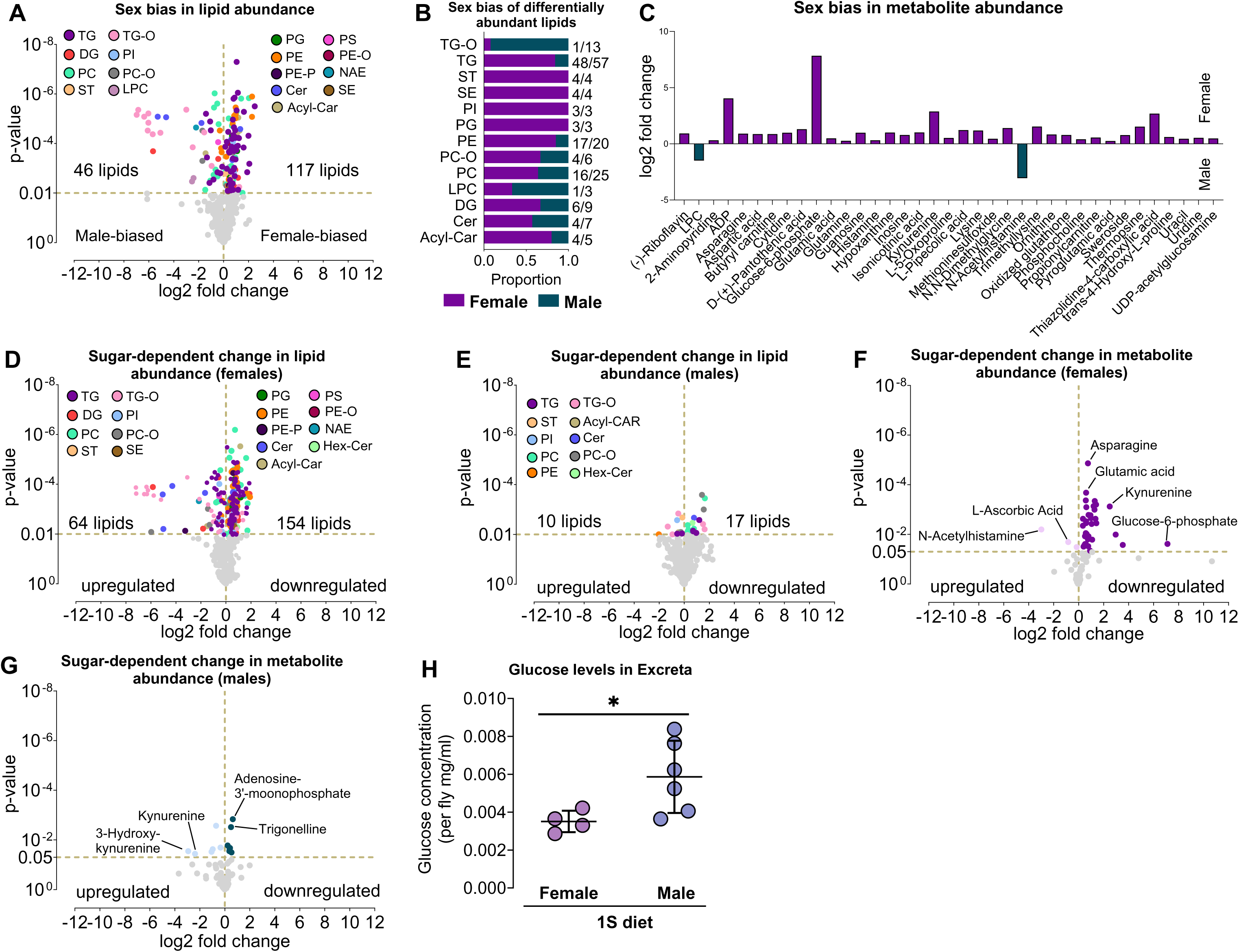
Female-biased effect of dietary sugar on metabolism. (A) Volcano plot of lipid species that are differentially regulated between 5-day-old *w^1118^* adult female and male fat bodies in 1S diet (colored dots represent *p*<0.01; Student’s *t*-test). (B) Number of lipid species within each sub-class that show higher abundance in either female (purple) or male (green) 5-day-old adult fat bodies in 1S diet (*p*<0.01; Student’s *t*-test). For example, 48/57 TG species are more abundant in females than in males, whereas 12/13 TG-O species are more abundant in males than in females. (C) Sex bias of all metabolites with differential abundance between female and male fat bodies in 1S diet (*p*<0.05; Student’s *t*-test). (D) Volcano plot of lipid species that are differentially regulated when females are maintained on a diet without sugar (colored dots represent *p*<0.01; Student’s *t*-test). (E) Volcano plot of lipid species that are differentially regulated when males are maintained on a diet without sugar (colored dots represent *p*<0.01; Student’s *t*-test). (F) Volcano plot of metabolites that are differentially regulated when females are maintained on a diet without sugar (colored dots represent *p*<0.05; Student’s *t*-test). (G) Volcano plot of metabolites that are differentially regulated when males are maintained on a diet without sugar (colored dots represent *p*<0.05; Student’s *t*-test). (H) Glucose levels in the excreta of 6-day-old adult *w^1118^* males were higher than females (*p*=0.0461, Student’s *t*-test); n=4-6 biological replicates. Data plotted as mean ± SEM. * *p*<0.05 TG: triglyceride; TG-O: ether-linked TG; DG: diglyceride; PI: phosphatidylinositol; PC: phosphatidylcholine; PC-O: ether-linked PC; ST: sterol; LPC: lyso-PC; PG: phosphatidylglycerol; PS: phosphatidylserine; PE: phosphatidylethanolamine; PE-O or PE-P: ether-linked PE; NAE: N-acetylethanolamine; Cer: Ceramide; Acyl-Car: acyl-carnitine; SE: sterol ester; Hex-Cer: hexosyl-ceramide. See also Figure S2.

When we examined specific fatty acids within triglyceride species that show a sex-bias, we found that triglycerides with a female bias contained a greater number of monounsaturated fatty acids such as myristoleic acid (C14:1), palmitoleic acid (C16:1), and oleic acid (C18:1) (Figure S2A). Triglycerides with a female bias also contained a greater number of odd-chain fatty acids (Figure S2A) and C28:2 (7/7). This female bias in odd-chain fatty acids aligns with published data showing odd-chain fatty acids are derived from the gut microbiome, where the microbe load is higher in adult female intestines than in males^88^. A female bias in C28:2 also aligns with the fact that it is a fatty acid precursor used for synthesis of female-specific pheromones. Triglycerides with a male bias contained a greater number of C24:1 (4/4) and myristic acid (C14:0) (Figure S2A). Beyond triglyceride, we also noted a significant female bias in lipid species within subclasses such as acyl-carnitine (acyl-Car) (4/5), ceramide (Cer) (4/7), diacylglycerol (DG) (6/9), phosphatidylcholine (PC) (16/25), ether-linked phosphatidylcholine (PC-O) (4/6), phosphatidylethanolamine (PE) (17/20), phosphatidylglycerol (PG) (3/3), phosphatidylinositol (PI) (3/3), sterol-ester (SE) (4/4) and sterol (ST) (4/4) (Figure 2B).

Lipid species with a significant male bias included lipids within subclasses such as lysophosphatidylcholine (LPC) (2/3) and ether-linked triacylglycerol (TG-O) (12/13) (Figure 2B). As with lipids, there was a significant female bias in many metabolites (*p*<0.05) (Figure 2C), including asparagine, glutamine, glutamic acid, glucose-6-phosphate, ADP, propionylcarnitine, pantothenic acid, which together suggest a female bias in glycolysis and the tricarboxylic acid (TCA) cycle. Supporting this, females showed a trend toward higher levels of citrate, a TCA cycle metabolite, than age- and genotype-matched males (*p*=0.06; Figure S2B). Thus, our data suggest that biological sex is a key regulator of the lipidome and metabolome.

To determine how dietary sugar affects the lipidome and metabolome in each sex, we compared these parameters in abdominal carcasses of flies transferred to the 1S or 0S diets as adults. In females, we observed significant differences in levels of 218 lipids between flies transferred to 1S compared with flies kept on 0S (*p*<0.01). Specifically, dietary sugar was associated with an increased abundance of 154 lipids and reduced abundance of 64 lipids (Figure 2D). Triglyceride showed the largest sugar-dependent change in abundance, where females maintained on 0S had significantly lower triglyceride abundance than females kept on 1S (61/81) (Figure S2C), consistent with our triglyceride assay data. When we examined specific fatty acids within triglyceride species affected by dietary sugar, we found that triglycerides with a bias toward females maintained on 0S had fewer C16:1 and C18:1 fatty acids (Figure S2D). Triglycerides with a bias toward females maintained on 0S similarly had fewer C28:2 fatty acids (Figure S2D). Broader effects of reduced dietary sugar were also observed across many lipid subclasses in females: Acyl-Car (3/4), Cer (5/9), DG (11/17), PC (29/33), PC-O (4/7), PE (25/27), PG (5/5), SE (1/1), ST (1/1), Hex-Cer (1/1), PE-O (1/1) (Figure S2C). Lipids that were significantly higher in abundance in females maintained on the 0S diet included NAE (1/1), PI (4/6), PS (1/1) and TG-O (17/21) (Figure S2C). Because body size is fixed before the adult stage of development^89^, and the females are only fed 1S or 0S after eclosion, these diet-induced alterations in lipid abundance cannot be explained by a sugar-dependent change in body size. In males, only 27 lipid species were differentially regulated by dietary sugar: 17 lipids were higher in abundance in flies maintained on a 1S diet whereas 10 lipids were lower in abundance (Figure 2E). Lipids that were lower in abundance in males on the 0S diet included PC (7/7), PC-O (2/3), triglyceride (4/6), Cer (1/1), Hex-Cer (1/1) (Figure S2E).

This female bias in sugar-induced changes in the fat body was reproduced in our metabolomic data (Supplemental Table S1, Figure 2F): 38 metabolites showed a difference in abundance in females maintained on the 0S diet whereas only 13 metabolites were altered in males kept on 0S (*p*<0.05 threshold for both sexes) (Figure 2G).

Metabolites that were altered in females on the 0S diet indicate reduced glycolysis and TCA cycle (asparagine, glutamine, glutamic acid, glucose-6-phosphate, ADP, nicotinamide, propionylcarnitine, pantothenic acid) and lower oxidative stress (oxidized glutathione, tans-4-hydroxy-L-proline, methioninesulfoxide, L-5-oxoproline (also known as pyroglutamic acid), kynurenine) (Figure 2F). In males, metabolites that were altered suggest reduced glycolysis on the 0S diet (nicotinamide, NAD, AMP) (Figure 2G). Using an independent assay to monitor sugar-dependent changes in TCA cycle activity, we found that both sexes had reduced citrate levels when maintained on 0S (Figure S2F). While metabolic flux analysis will be an important future direction to extend knowledge of sex differences in the metabolic fate of dietary sugar, our data suggest sugar caused a strongly female-biased remodeling of the lipidome and metabolome in abdominal carcasses. While the reason for this profound female bias in changes to the lipidome and metabolome in response to dietary sugar is unclear, males showed higher levels of sugar excretion on the 1S diet (Figure 2H).

### Dietary sugar promotes sex differences in IIS regulation

Considering dietary sugar stimulates the production and secretion of Dilps from the IPCs in the brain^90–93^, we tested if Dilp production, Dilp secretion, and IIS activity differed between males and females maintained on 1S. We first measured mRNA levels of IPC-derived *dilp2*, *dilp3* and *dilp5* in adult heads using quantitative real-time polymerase chain reaction (qPCR). Female heads had higher mRNA levels of *dilp3* than male heads (Figure 3A); no sex differences in mRNA levels of *dilp2* and *dilp5* were found (Figure 3B and 3C). To test whether dietary sugar caused the sex difference in *dilp3* mRNA levels, we compared *dilp3* mRNA levels in males and females transferred to either 1S or 0S diets as adults. Female flies transferred to 0S had significantly lower *dilp3* mRNA levels than females kept on 1S (Figure 3D). Because there was no sugar-dependent change in male *dilp3* levels, the sex difference in *dilp3* levels was eliminated (Figure 3D; sex:diet interaction *p*=0.0128), showing dietary sugar plays a role in the female bias in *dilp3* mRNA levels. We next asked whether this bias in *dilp3* mRNA levels extended to Dilp3 protein using a published anti-Dilp3 antibody^94^. We observed a female bias in Dilp3 levels in the IPC of 5-day-old adults in both the 1S and 0S diets (Figure S3A-S3E), confirming Dilp3 levels are higher in females than in males.

**Figure 3.**
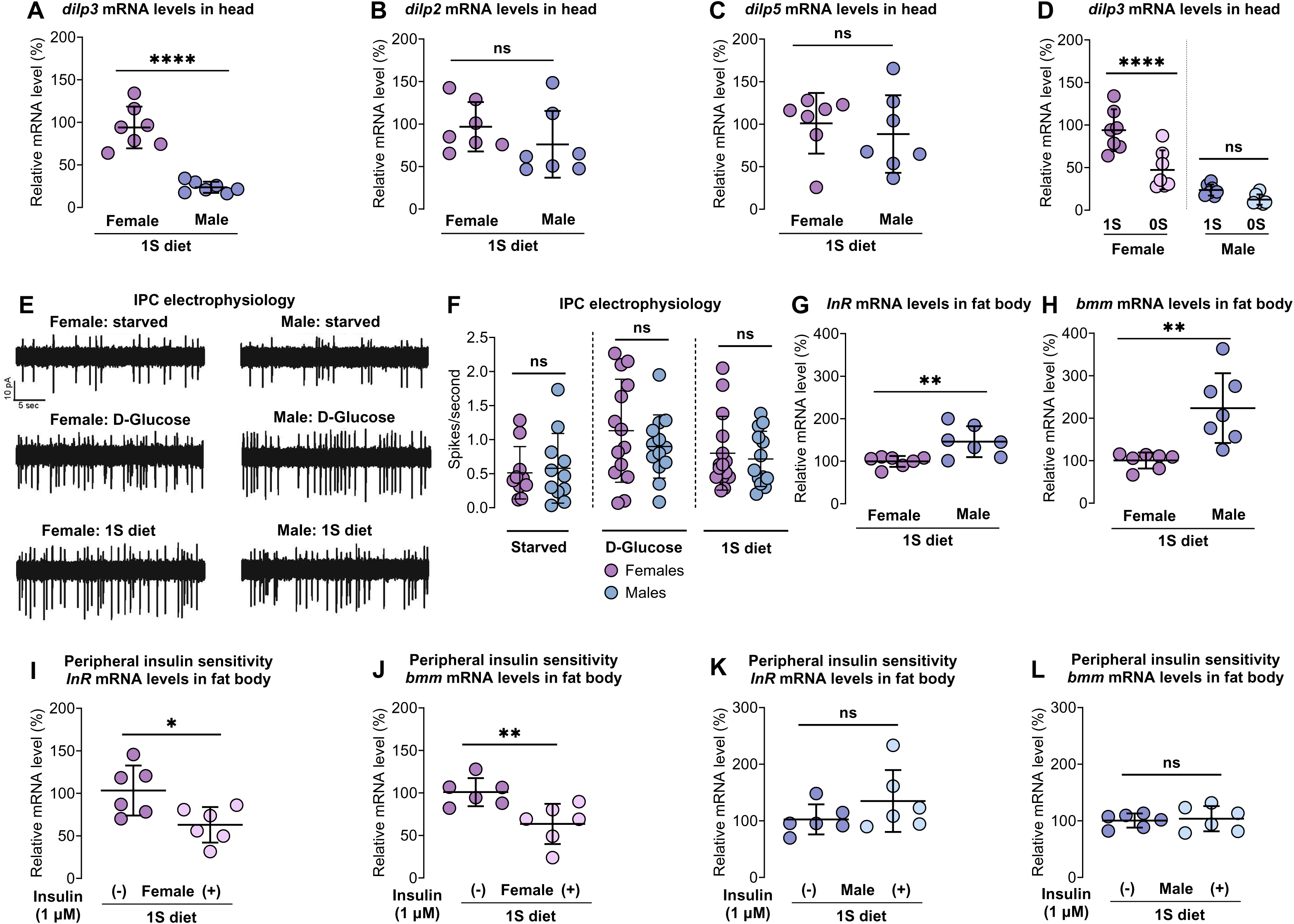
Dietary sugar promotes sex differences in IIS regulation. (A) mRNA levels of IPC-derived *dilp3* in heads were significantly higher in 5-day-old *w^1118^* females compared with males (*p*<0.0001; Student’s *t*-test); n=7 biological replicates. (B, C) mRNA levels of IPC-derived *dilp2* (B) and *dilp5* (C) in heads were not significantly different between 5-day-old *w^1118^* males and females (*p*=0.1282, Mann-Whitney test (*dilp2*); *p*=0.5753, Student’s *t*-test (*dilp5*)); n=7 biological replicates. (D) mRNA levels of IPC-derived *dilp3* in heads were significantly lower in females maintained on the 0S diet compared to females kept on the 1S diet (*p*<0.0001). There was no sugar-dependent change in *dilp3* mRNA in males (*p*=0.4809; sex:diet interaction *p*=0.0128). Two-way ANOVA followed by Bonferroni post-hoc test; n=7 biological replicates. Note: *dilp3* mRNA levels in 1S diet were replotted from Figure 3A as these data were collected in the same experiment. (E) Representative cell-attached recordings of IPC activity in 4-8-day-old female (left) and male (right) flies that were either starved for 24 hr (top), re-fed with D-glucose (middle), or re-fed with whole food (bottom). (F) Quantification of cell-attached recordings of IPC activity in 4-8-day-old male and female flies that were either starved for 24 hr, re-fed with D-glucose, or re-fed with whole food. n=10-15 individuals per sex and per condition; error bars indicate SEM. (G, H) mRNA levels of Foxo target genes-*InR* (G) *and bmm* (H) in fat bodies were significantly higher in 5-day-old *w^1118^* males compared to females in 1S diet (*p*=0.0077 (*InR*); *p*=0.0022 (*bmm*) Student’s *t*-test); n=7 biological replicates. (I, J) mRNA levels of Foxo target genes-*InR* (I) and *bmm* (J) in 5-day-old *w^1118^* female fat bodies were significantly reduced after stimulation with 1 µM insulin compared with fat bodies lacking insulin stimulation (*p*=0.0208 (*InR*); *p*=0.0100 (*bmm*); Student’s *t*-test); n=6 biological replicates. (K, L) mRNA levels of Foxo target genes-*InR* (K) and *bmm* (L) in 5-day-old *w^1118^* male fat bodies were not significantly different between unstimulated and insulin-stimulated conditions (*p*=0.2194 (*InR*); *p=*0.7649 (*bmm*); Student’s *t*-test); n=6 biological replicates. Data plotted as mean ± SEM. ns indicates not significant with *p*>0.05; * *p*<0.05, ** *p*<0.01, *** *p*<0.001, **** *p*<0.0001. See also Figure S3.

Given the sex bias in *dilp3* mRNA and protein, we next monitored Dilp secretion from the IPCs. We first expressed a calcium-responsive chimeric transcription factor LexA-VP16-NFAT (*UAS-LexA-VP16-NFAT* [called *UAS-CaLexA*]) in the IPC using *dilp2-GAL4* (full genotype *dilp2-GAL4>UAS-LexA-VP16-NFAT*). Because the chimeric transcription factor drives expression of GFP in response to changes in cellular calcium levels, this method provides an indirect readout of neuronal activity and Dilp release^95^. On the 1S diet, there was no sex difference in GFP expression within the IPC (Figure S3F-S3H). To confirm this, we directly monitored IPC activity using cell-attached electrophysiological recordings. When we examined IPC spike rates, we found no significant differences between males and females after a period of fasting (Figure 3E and 3F), or when the fasting period was followed with refeeding on either whole food or D-glucose (Figure 3E and 3F). These data suggest IPC activity is equivalent between the sexes, confirming results from another recent study^96^. Given that Dilp3 levels in the IPC show a female bias, these data suggest females have a greater quantity of Dilp3 available for secretion each time the IPC are active.

To test whether the female bias in *dilp3* mRNA caused by dietary sugar was due to an intrinsic difference in IPC, we expressed sex determination gene *transformer* (*tra*) in male IPC and monitored *dilp3* mRNA levels. Normally, a functional Tra protein is expressed only in females, where it directs most aspects of female sexual development^97–100^. Because Tra expression in males specifies female development across multiple tissues^101–107^, this allows a female sexual identity to be conferred upon male IPC. In males with IPC-specific expression of Tra, we observed no significant change in *dilp3* mRNA levels (Figure S3I). This suggests the sexual identity of the IPC does not determine the sex difference in *dilp3* mRNA levels. While the APCs have also been shown to influence *dilp3* mRNA levels in larvae^108^, Tra expression in these cells similarly had no effect on the sex difference in *dilp3* mRNA in adults (Figure S3J). Given that many groups of neurons and neuropeptide-producing cells have been shown to influence *dilp3* mRNA levels in a non-cell-autonomous manner^43,62, 109^, we next used *elav-GAL4* to drive Tra expression broadly across these cells. We found that Tra expression in all *elav-GAL4*-positive cells caused a significant increase in male *dilp3* mRNA levels (Figure S3K), suggesting that Tra acts in neurons or neuropeptide-producing cells other than the IPC to regulate the sex difference in *dilp3* mRNA levels.

Given the female bias in *dilp3* mRNA and protein, and the known role of *dilp3* in promoting IIS activity^108,110,111^, we next compared IIS activity in males and females. To assess IIS activity, we measured mRNA from known transcriptional targets of Forkhead box, sub-group O (Foxo; FBgn0038197) in male and female abdominal carcasses. When IIS activity is high, Foxo is phosphorylated and repressed, leading to lower mRNA levels of known target genes *InR* and *bmm*^112–115^. In adult males and females maintained on 1S, mRNA levels of Foxo target genes were significantly lower in females than in males (Figure 3G and 3H). This suggests IIS activity is normally higher in females than in males. Given that *Drosophila* female larvae are more insulin sensitive than male larvae^80^, a sex difference that also exists across many tissues in mammals^116,117^, we also compared insulin sensitivity between adult virgin males and females kept on 1S. Considering female abdominal carcasses showed significant insulin-induced repression of Foxo target genes (Figure 3I and 3J), a response we did not observe in males (Figure 3K and 3L), our data suggest female abdominal carcasses are more insulin sensitive than males. Taken together, our data show that dietary sugar promotes higher expression of *dilp3* in females. Increased *dilp3* mRNA and protein, combined with higher insulin sensitivity in females, likely contribute to elevated IIS activity in females maintained on 1S.

### Females require IIS to maintain fat storage

Insulin is a potent lipogenic hormone in multiple animals including *Drosophila*^53,118–122^. Increased IIS activity in female abdominal carcasses may therefore explain their higher fat storage. Because reduced IIS activity during development significantly impairs larval growth and survival to adulthood^54,103,110,111,123–125^, and increased IIS activity during development augments body size^103,125–127^, we needed a system to modulate IIS activity specifically in adults. Unfortunately, existing systems for inducible gene expression have undesirable limitations for studying sex differences in fat metabolism. For example, the Gene Switch system^128,129^ induces transgene expression by feeding flies mifepristone, which has sex-specific effects on adult physiology^130–132^. The AGES (Auxin-inducible Gene Expression System) method uses auxin feeding to induce transgene expression^133^, which we and others recently showed affects feeding and triglyceride metabolism in a sex-biased manner^134^.

Another inducible system for gene expression, the temporal and regional gene expression targeting (TARGET) system^135^, normally requires long-term temperature shifts at 29°C to inactivate temperature-sensitive repressor GAL80. Given that inactive GAL80 cannot repress GAL4, UAS transgene-mediated expression is initiated as long as the animals are kept at 29°C. Unfortunately, 29°C is a temperature known to disrupt lipid metabolism^136,137^, making a long-term heat shock to induce transgene expression or RNAi-mediated knockdown undesirable. To overcome this limitation, we reasoned that we could deliver a brief 3 hr heat shock to newly-eclosed male and female flies to transiently express proapoptotic gene *reaper* (*rpr*) specifically in the IPC (genotype: *dilp2-GAL4>UAS-rpr,tub-GAL80^ts^*). To validate expression of *dilp2-GAL4* in each sex, we first confirmed that *dilp2-GAL4*-positive cells were morphologically similar to published data^45,138–140^, with no obvious differences between males and females (Figure S4A and S4B). We also confirmed *dilp2-GAL4*-positive cells express Dilp3 (Figure S4C, S4CL, S4CLL, S4D, S4DL, S4DLL), in line with published data^43,138^. We next assessed whether *dilp2-GAL4* could be used to ablate the IPC. Importantly, mediated *rpr* overexpression is an established way to ablate the IPC^123,141,142^. Based on published literature^110,111,123,142,143^, in adult heads the IPC are the only place that *dilp2* and *dilp3* mRNA are produced.

Therefore, we measured *dilp2* and *dilp3* mRNA levels as a direct way to test IPC ablation.

Supporting successful IPC ablation, mRNA levels of *dilp3* and *dilp2* were reduced in adult heads from female and male flies after a brief heat shock compared with genotype-matched controls without heat shock (Figure 4A, 4B, S4E, S4F). When we used anti-Dilp3 to check IPC ablation, many flies lacked Dilp3-positive cells in the *pars intercerebralis* region of the brain where the IPC are located, indicating a complete IPC ablation in these individuals (Figure S4G-S4J). In line with prior reports^141,142^, some individuals had residual Dilp3-producing cells, suggesting a partial IPC ablation (Figure S4K and S4L). Given that our approach to fully/partially ablate the IPC caused phenotypes associated with IPC loss and reduced insulin signaling such as an extremely long lifespan in both sexes (Figure 4C and 4D)^141,144–148^ and transiently reduced fertility in female but not male flies (Figure 4E and 4F)^142^, we conclude that our approach to IPC ablation is a valid method for assessing how adult-specific reduction in IIS affects fat storage. While the reason for the transient effect on female fertility remains unclear, previous studies show that changes to insulin affect female pheromones^149^ which could introduce a minor delay in egg-laying.

**Figure 4.**
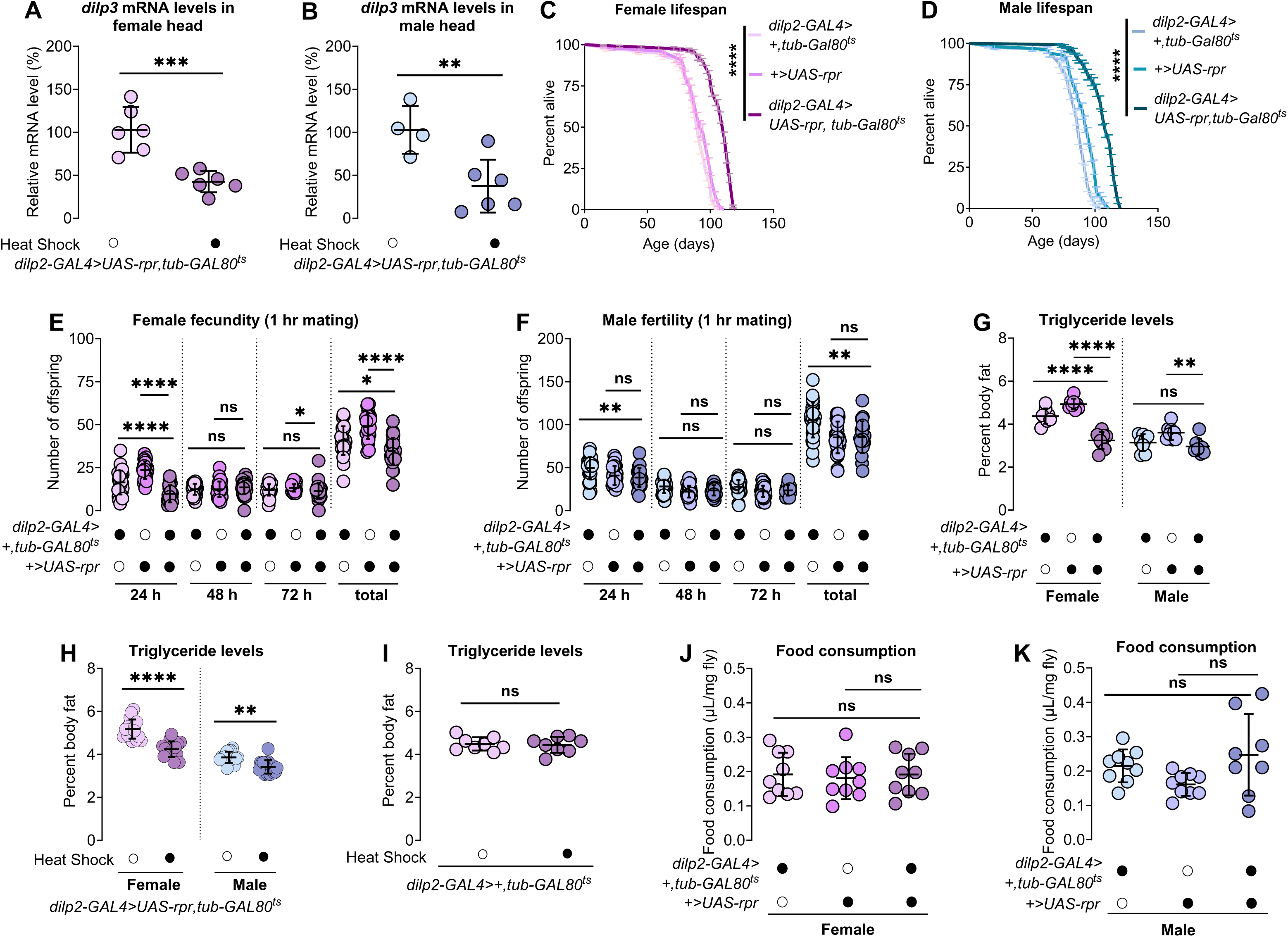
Females require IIS to maintain fat storage. (A, B) mRNA levels of IPC-derived *dilp3* in heads were significantly lower in both female (A; *p=*0.0005, Student’s *t*-test) and male (B; *p*=0.0092, Student’s *t*-test) *dilp2-GAL4>UAS-rpr,tub-GAL80^ts^* flies with heat shock at 29°C compared to sex-matched flies reared in room temperature; n=4-6 biological replicates. (C) Lifespan was significantly longer in *dilp2-GAL4>UAS-rpr,tub-GAL80^ts^* females compared with *dilp2-GAL4>+,tub-GAL80^ts^* and +>*UAS-rpr* controls (*p*<0.0001 and *p*<0.0001, respectively; Log-rank test); n=100-213 females. (D) Lifespan was significantly longer in *dilp2-GAL4>UAS-rpr,tub-GAL80^ts^*males compared with *dilp2-GAL4>+,tub-GAL80^ts^* and +>*UAS-rpr* controls (*p*<0.0001 and *p*<0.0001, respectively; Log-rank test); n=100-148 males. (E) The number of eggs laid by females in the first 24-hour period post-mating that developed into pupae was significantly lower in *dilp2-GAL4>UAS-rpr,tub-GAL80^ts^*females compared with both *dilp2-GAL4>+,tub-GAL80^ts^* and +>*UAS-rpr* controls (*p*<0.0001 and *p*<0.0001, respectively; one-way ANOVA followed by Tukey HSD). n=30 females. (F) The number of eggs laid by females mated to *dilp2-GAL4>UAS-rpr; tub-GAL80^ts^* males were significantly lower compared to control *dilp2-GAL4>+; tub-GAL80^ts^* but no change compared to +>*UAS-rpr* control males that developed into pupae over the first 24 hr of after 24 hr mating (*p*=0.0032 and *p*=0.7420, respectively; one-way ANOVA followed by Tukey HSD). n=22-30 males. (G) Whole-body triglyceride levels were significantly lower in *dilp2-GAL4>UAS-rpr,tub-GAL80^ts^* females compared with both *dilp2-GAL4>+,tub-GAL80^ts^* and *+>UAS-rpr* control females (*p*<0.0001 and *p*<0.0001, respectively). Whole-body triglyceride levels were not significantly different in *dilp2-GAL4>UAS-rpr,tub-GAL80^ts^* males compared with *dilp2-GAL4>+,tub-GAL80^ts^* control males but significantly different from *+>UAS-rpr* control males (*p*=0.9397 and *p*=0.0022, respectively) (sex:genotype interaction *p*=0.0002). Two-way ANOVA followed by Bonferroni post-hoc test; n=8 biological replicates. (H) Whole-body triglyceride levels were significantly lower in female *dilp2-GAL4>UAS-rpr,tub-GAL80^ts^* after heat shock compared with genotype-matched females without heat shock (*p*<0.0001). A reduction in whole-body triglyceride levels was also observed in males (*p*=0.0012); however, the magnitude of the decrease was smaller than in females (sex:genotype interaction *p*=0.0071). Two-way ANOVA followed by Bonferroni post-hoc test; n=16 biological replicates. (I) Whole-body triglyceride levels were not significantly different between *dilp2-GAL4>+,tub-GAL80^ts^* females that received a brief heat shock compared with *dilp2-GAL4>+,tub-GAL80^ts^* females that received no heat shock (*p*=0.8107; Student’s *t*-test); n=8 biological replicates. (J, K) Food consumption was not significantly different in either females (J) or males (K) between *dilp2-GAL4>UAS-rpr,tub-GAL80^ts^*flies and *dilp2-GAL4>+,tub-GAL80^ts^* or *+>UAS-rpr* controls (females: *p*=0.9999 [GAL4 control] and *p*=0.9318 [UAS control]; males: *p*=0.6516 [GAL4 control] and *p*=0.0628 [UAS control]). One-way ANOVA followed by Tukey HSD; n=8-9 biological replicates. All data in (A, B, E-K) plotted as mean ± SEM. ns indicates not significant with *p*>0.05, \**p*<0.05, ** *p*<0.01, *** *p*<0.001, **** *p*<0.0001. See also Figure S4.

Using this system to ablate the IPC, we tested whether increased IIS was required for females to store more body fat than males. We found that triglyceride levels were significantly lower in 5-day-old females with ablated IPCs (genotype *dilp2-GAL4>UAS-rpr,tub-GAL80^ts^*) than in control females (genotypes *dilp2-GAL4>+,tub-GAL80^ts^* and +>*UAS-rpr*) (Figure 4G and S4M). In contrast, IPC ablation in males had no significant effect on body fat (Figure 4G and S4M). We also found that females (genotype *dilp2-GAL4>UAS-rpr,tub-GAL80^ts^*) subjected to a brief heat shock had lower body fat compared with genotype-matched females that were not heat-shocked (Figure 4H). While heat-shocked *dilp2-GAL4>UAS-rpr,tub-GAL80^ts^* males showed a decrease in body fat (Figure 4H), the magnitude of the decrease was greater in females (sex:genotype interaction *p*=0.0071) (Figure S4N). The reduction in female body fat upon adult-specific loss of IPC was not explained by the brief heat shocks, as control females (genotype *dilp2-GAL4>+,tub-GAL80^ts^*) subjected to an identical heat shock showed no significant effect on body fat (Figure 4I).

The decrease in body fat was also not due to altered feeding in animals with IPC ablation, as no significant differences in food consumption were present in either females or males with IPC ablation compared with controls (Figure 4J and 4K). Instead, we identify *dilp3* as a key factor that contributes to higher female fat storage when dietary sugar is present: Dilp3 overexpression in the IPC blocked the decrease in body fat normally observed in adult females maintained on 0S (Figure S4O; diet:genotype interaction *p*<0.0001). Dilp3 overexpression had no effect on sugar-induced changes to body fat in males (diet:genotype interaction *p*=0.5588), possibly due to insulin resistance (Figure S4P). Supporting a specific role for IIS in maintaining higher fat storage in females, supplementing the diet of virgin females with an ecdysteroid called 20-hydroxyecdysone (20HE) did not promote fat storage in females (Figure S4Q). Our data therefore reveal sex differences in IIS regulation in adult flies, and demonstrate the importance of this regulation to the male-female difference in fat storage.

## Discussion

Our findings provide significant insight into the mechanisms by which adult females store more fat than males. This information is critical to our understanding of how sex differences in body fat are established and maintained, as past studies identified mechanisms that operate in males with little effect in females^5,10^. Based on our data, we propose a model in which higher consumption of dietary sugar in females leads to elevated *dilp3* mRNA and protein levels, which act together with greater insulin sensitivity to promote higher IIS activity and body fat in females. Supporting this model, reduced sugar consumption or IPC ablation lessened the sex difference in fat storage by lowering body fat in females with little effect in males, and Dilp3 overexpression prevented the reduction in fat storage normally observed in females maintained on 0S. This reveals previously unrecognized sex-specific IIS regulation in adult flies, and identifies Dilp3 and IIS as key factors that support increased fat storage in adult females.

One important advance in our study was to extend knowledge of how food intake affects physiology in unmated females. Past studies used mated females to show that greater food consumption supports the quality and quantity of eggs produced^9,150–157^. While greater total nutrient intake could explain these effects, specific macronutrients such as dietary protein play individual roles in supporting egg production, egg-laying, and postcopulatory behaviors in mated females^158–160^. Indeed, dietary protein is limiting for *Drosophila* egg production^161^, where its biological importance is shown by the fact that mated females will adjust their food preferences to ensure adequate protein intake^159,162,163^. Dietary protein does not hold the same physiological value for unmated females, however, as virgin females do not adjust their preferences after protein restriction^163^. Instead, our data suggest that dietary sugar is a key macronutrient for virgin female physiology due to its effects on IIS activity and fat accumulation. Because IIS activity supports normal germline stem cell proliferation and maintenance in the ovary^54,142,161,164–167^, and high levels of body fat are required for successful egg production^12^, dietary sugar consumption likely helps to establish a supportive metabolic state prior to reproduction. Once mated, this role for dietary sugar becomes less important, as reflected by the limited consumption of a sugar-rich diet in mated females compared with males^168^. Thus, our data suggests that higher food intake in both virgin and mated females provide key macronutrients to fuel the specific needs that accompany particular life history stages (*e.g*. prior to mating or during egg production).

Another advance of our study was to reveal a profound sex difference in how dietary sugar affects metabolism in the fat body. In females, sugar caused profound remodeling of the lipidome and many changes in the metabolome. This sex difference was most apparent when examining triglyceride, the lipid sub-class with the most obvious sugar-dependent changes in female lipid abundance. However, females showed a sugar-dependent increase across many lipid sub-classes (*e.g*., DG, PG, PE, Acyl-Car). Given that females have lower lipase activity than males^5^, the presence of high diglyceride and triglyceride levels suggest increased lipid synthesis in females. Diglyceride is also a precursor for phospholipids such as PE and PC synthesis^1,169^. High levels of PE and PC in females maintained on 1S further support higher lipid synthesis in females, a possibility that will be important to test in future studies.

Beyond this broad lipid accumulation in females kept on 1S, other interesting findings were that dietary sugar enhanced levels of a fatty acid precursor used for synthesis of female-specific pheromones (28:2) and odd-chain fatty acids in the triglyceride pool. Fatty acid 28:2 is the direct precursor for the synthesis of a cuticular hydrocarbon called 7,11-heptacosadiene, the major female pheromone that mediates female sexual attractiveness^170–172^. Thus, in addition to supporting IIS activity and fat accumulation in females, dietary sugar consumption may also ensure females emit an optimal pheromone profile. Indeed, sexual attractiveness is altered in females with abnormal levels of cuticular hydrocarbons^173^. Given that the gut microbiota help produce odd-chain fatty acids^174^ and the microbe load is higher in adult female intestines than in males^88^, our data suggest that the intestinal microbiome may provide precursors used for fat storage in females exposed to dietary sugar. Thus, dietary sugar provides an array of physiological benefits for females to optimize reproduction.

In males, dietary sugar caused few changes in either lipids or metabolites. One notable finding was that only one lipid subclass, TG-O, showed higher abundance in males than in females. Ether-linked lipids such as TG-O are synthesized by a multi-step process mediated by enzymes located both in the peroxisome and endoplasmic reticulum^175–177^.

Because a recent study found high levels of TG-O in male flies chronically fed a high-sugar diet^178^, higher TG-O levels in males may indicate sugar-induced lipotoxicity. Indeed, elevated TG-O levels were associated with metabolic dysfunction, as genetically reducing production of ether-linked lipids in male flies reversed defects in glucose levels, cardiac function, and insulin signaling caused by the high-sugar diet^178,179^. Future studies will therefore need to examine ether-linked lipid synthesis in both sexes to determine whether male-female differences exist in this process and its relationship to metabolic dysfunction. Follow-up studies will also need to determine the ultimate reason that males largely fail to adjust the lipidome in response to dietary sugar. Potential explanations suggested by our data include insulin resistance, increased sugar excretion, and high lipase and lipolytic pathway activity^5,10^. Published data suggest that greater carbohydrate breakdown within the male gut to support spermatogenesis might also explain the female-biased lipidomic response to dietary sugar^105^.

Other key findings of our study include showing that dietary sugar promotes a female bias in *dilp3* levels, and revealing a female bias in IIS activity within the adult fat body. This extends knowledge from past studies on sex differences in IIS activity in third-instar larvae^103^ by showing this female bias in IIS activity is present in adults and is physiologically significant. Given that elevated *dilp3* levels are known to promote body fat^43,62,180^ and prolong starvation resistance^61,181,182^, and that Dilp3 overexpression blocks the reduction in body fat normally observed on the 0S diet, this female bias in *dilp3* is significant for higher fat accumulation in females. However, a number of important future directions remain. First, it will be important to determine whether Dilp3 signaling via the insulin receptor drives all the lipidomic changes we observed due to dietary sugar consumption, or whether other Dilps play a role. Second, it will be important in future studies to reveal the mechanisms underlying the sex bias in *dilp3* levels. We identified sex determination gene Tra as a key regulator of the sex difference in *dilp3* mRNA levels.

While it is clear that Tra does not act directly on the IPC or in the Akh-producing cells to influence the sex difference in *dilp3* mRNA levels, the cell type in which Tra acts to specify this difference remains unclear. Future studies will need to determine whether Tra acts in cells that produce *dilp3*-regulatory factors such as gut-derived neuropeptide F^43^, octopamine^181^, tachykinin^109^, leucokinin^61,183^, adiponectin receptor^184^, Allatostatin A^45^, bursicon^62^, drosulfakinin^182^, and potentially Akh^108^ and Limostatin^180,185^ to regulate the sex difference in *dilp3* mRNA levels, and whether this involves changes in food intake. Indeed, sex differences in expression of select *dilp3*-regulatory factors have been reported^10,104,186^, where these factors have sex-specific effects on physiology^10,42–46,61,62,109,180,181^ and influence mating and/or reproductive behaviors^187–195^.

Overall, our findings identify IIS as a major determinant of female fat storage, with only minor effects on male body fat. This contrasts with males, where a set point of stored fat is maintained by *bmm* and Akh, pathways with little effect on female body fat^5,10^. The primary mechanisms that establish fat storage are therefore not identical between the sexes: females sustain body fat primarily via the anabolic effects of insulin whereas body fat in males is maintained by the lipolytic effects of *bmm* and Akh. Future studies will need to determine whether the sex-biased effects of changing IIS and Akh on body fat are reproduced in other aspects of physiology regulated by these pathways (*e.g*. locomotion, lifespan, aggression, feeding, sleep)^64,141,142,149,152,196–199^. Future studies will also need to determine whether the extensive crosstalk between IIS and Akh identified in single-and mixed-sex studies in adults and larvae is shared between males and females^45,57,108,123,200^. Because IIS and Akh mediate responses to a variety of extrinsic and intrinsic cues, gaining a better understanding of the interplay between these two major hormone pathways in males and females will provide critical insights into how diverse cues are integrated to maintain whole-body triglyceride homeostasis in each sex. Given that sex differences in sucrose-induced changes to mammalian adipose tissue lipolysis, hepatic lipogenesis, and adipose tissue gene expression broadly reproduce the effects we report in flies^201^, the sex-biased effects of IIS and Akh on body fat we observed have implications for our understanding of sex differences in parallel hormone systems in mammals (*e.g*. insulin, glucagon) and the sex-biased risk of diseases associated with these hormones^202–206^.

### Limitations of the study

Our study has several limitations. First, the limitations of current *Drosophila* inducible gene expression systems mean that we overexpressed *dilp3* throughout development instead of just in adults. We therefore cannot rule out the possibility that Dilp3 acts in development to influence adult fat storage. Second, while we defined a role for Dilp3 in regulating the sex difference in body fat, we cannot rule out a role for other Dilps.

### Resource availability

#### Lead contact

Further information and requests for resources and reagents should be directed to, and will be fulfilled by, lead contact Elizabeth J. Rideout.

#### Materials availability

This study did not generate new unique reagents.

## Data and code availability

- Fly lipidomic and metabolomic data are available in Supplemental Table S1.
- This paper does not report original code.
- Any additional information required to reanalyze the data reported in this paper is available from the lead contact upon request.

## Acknowledgments

This study was supported by operating grants to EJR from the Canadian Institutes for Health Research (PJT-153072 and PJT-183786), CIHR Sex and Gender Science chair program (GS4-171365), Michael Smith Foundation for Health Research (16876), and the Canadian Foundation for Innovation (JELF-34879). PB was supported by a 4-year CELL fellowship from UBC. LWW was supported by a British Columbia Graduate Scholarship Award. TH/HY were supported by Natural Sciences and Engineering Research Council of Canada (NSERC; 2020-04895). MDG and MS were supported by NSERC (RGPIN-2016-03857). We used FlyBase to find information, stocks, and literature for our project. FlyBase is supported by a grant from the National Human Genome Research Institute at the U.S. National Institutes of Health (U41 HG000739) and by the British Medical Research Council (MR/N030117/1). Stocks obtained from the Bloomington *Drosophila* Stock Center (NIHP40OD018537) were used in this study. The authors thank the TRiP at Harvard Medical School (NIH/NIGMS R01-GM084947) for providing transgenic RNAi fly stocks and/or plasmid vectors used in this study. We also thank Dr. Jan Veenstra (University of Bordeaux) and Dr. Patrick Callaerts (University of Leuven) for providing anti-Dilp3 antibody, and Dr. Gaiti Hasan (National Centre for Biological Science, India) for providing *UAS-dilp3* flies. The graphical abstract was prepared using BioRender (https://BioRender.com). The funders had no role in study design, data collection and analysis, decision to publish, or preparation of the manuscript. We thank members of the Rideout lab for valuable feedback. We acknowledge that our research takes place on the traditional, ancestral, and unceded territory of the Musqueam people; a privilege for which we are grateful.

## Author contributions

P. B. conceived studies, collected samples, conducted experiments, interpreted experiments, wrote and edited the manuscript.

J. A. B. and J. B. L. performed dissections and collected data related to brain anti-Dilp3 immunofluorescence.

H. Y. performed lipidomic and metabolomic experiments.

L. W. W. performed ecdysone experiment.

C. J. M. performed brain dissections and monitored IPC activity using the CaLexA system.

M. D. G. edited the manuscript and acquired funding for electrophysiological recordings of IPC.

T. H. edited the manuscript and acquired funding for lipidomic and metabolomic analyses.

M. S. conceived studies, analyzed data, interpreted experiments, and contributed to writing and editing of the manuscript related to electrophysiological recordings of IPC.

E. J. R. conceived studies, interpreted experiments, wrote and edited the manuscript, acquired funding, and is the guarantor of this work.

## Supplemental information

Document S1. Supplemental Figures S1-S4

Table S1. Excel file containing raw data with calculations

Table S2. Excel file containing statistics for all data

## STAR⍰Methods

### Key resources table

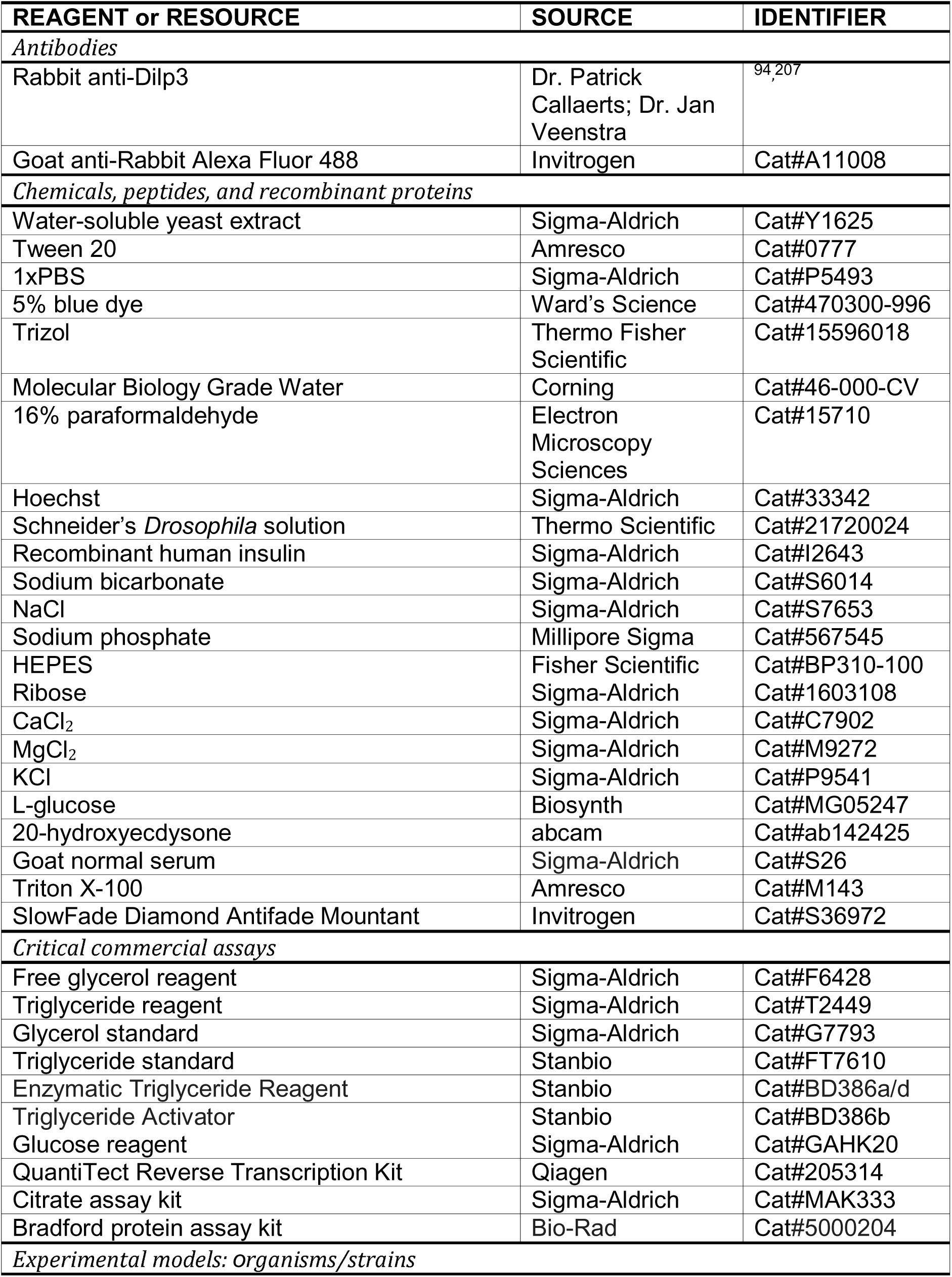
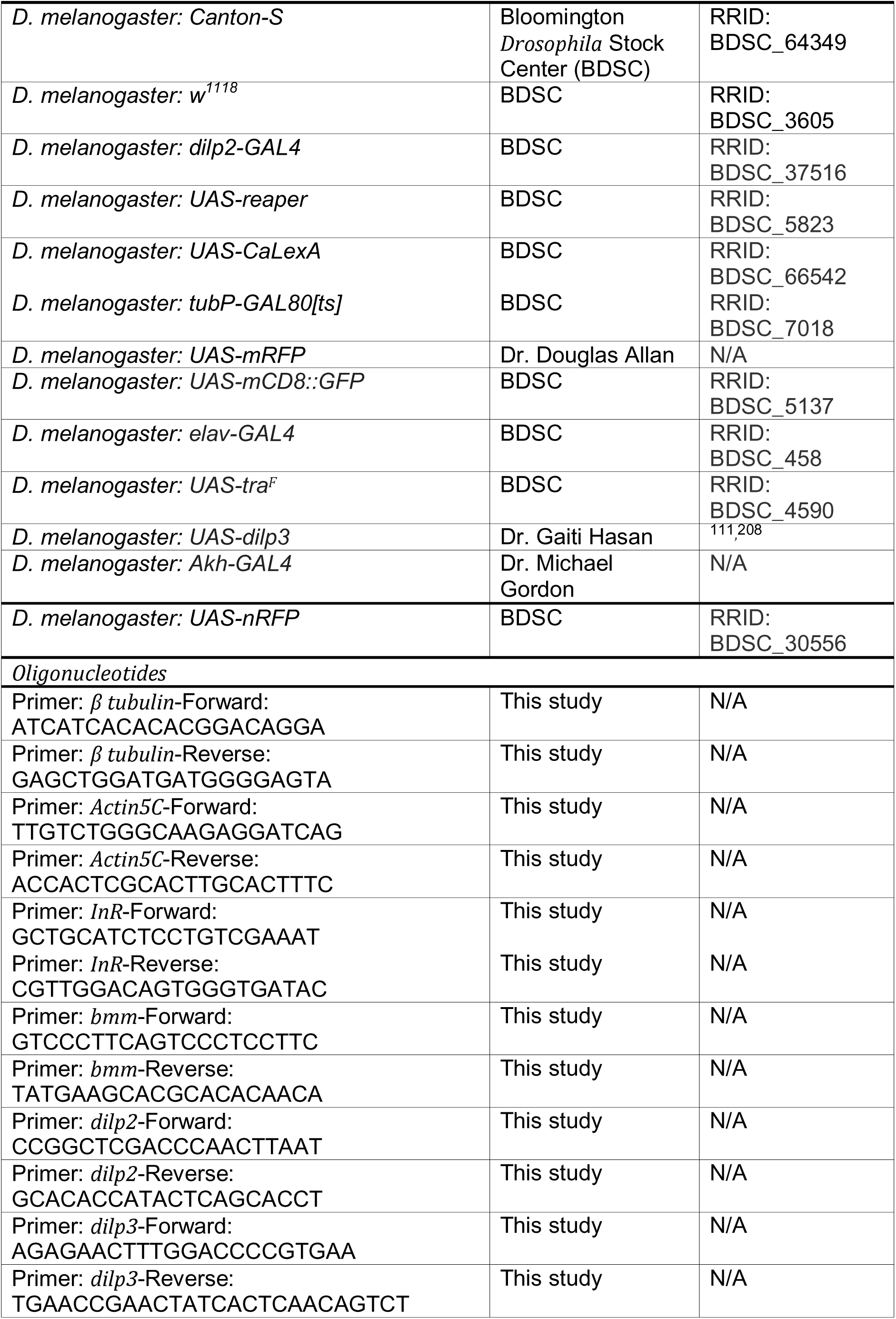
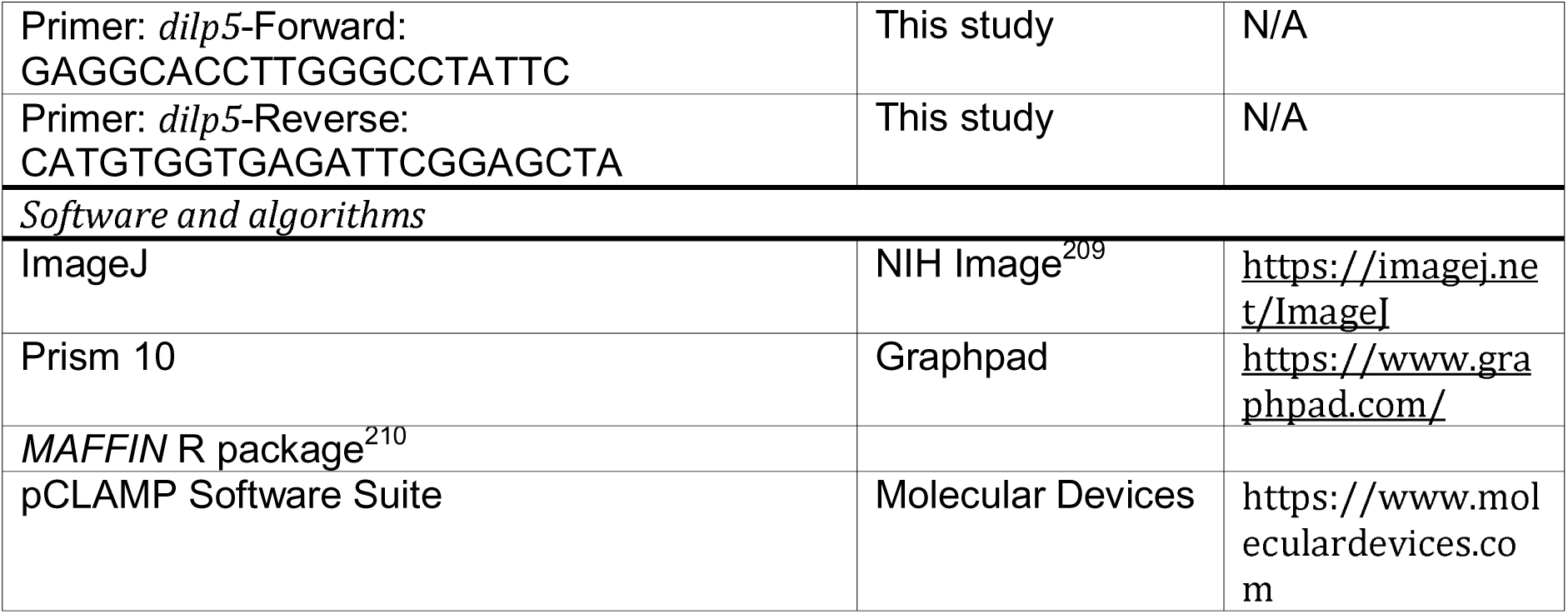

### Experimental model and study participant details Fly husbandry

Detailed lists of strains are provided in the key resources table. The recipe for our regular lab diet (1S) is as follows: 20.5 g/L sucrose, 70.9 g/L D-glucose, 48.5 g/L cornmeal, 45.3 g/L yeast, 4.55 g/L agar, 0.5g CaCl2•2H2O, 0.5 g MgSO4•7H2O, 11.77 mL/L acid mix (propionic acid/phosphoric acid). To make 1 L of 20% food, 800 mL 0.7% agar was mixed with 200 mL of 1S food. The low-sugar diet (0S) was made by leaving out sucrose and D-glucose from the 1S diet. The 1S diet was calorie-matched to 0S (1S-low calorie) by removing 8.6 g/L cornmeal from the 1S diet. For diets with sugar added back, either 20.5 g/L sucrose (0S+sucrose) or 70.9 g/L D-glucose (0S+D-glucose) were added to 0S, as this reflects the usual quantity of each sugar in the 1S diet. To monitor the effect of non-nutritive sugar on body fat, we added 62.5 g/L L-glucose (0S+L-glucose; Biosynth, MG05247) to the 0S diet^87^. To make a 30% high sugar diet, 300 g/L extra sucrose was added to the 1S diet, in line with published work^211,212^. For ecdysone-supplemented food, 0.5 mg/mL of 20-hydroxyecdysone (20-HE) in ethanol was added to the food (abcam, ab142425). For control food, the same ethanol volume without 20-HE was added to the food. For all experiments, we allowed female flies to lay eggs on grape juice agar plates for 12 hr. At 24 hr after egg laying, 50 larvae were picked into vials containing 10 mL food and reared at 22°C. Males and females were distinguished by the presence of sex combs late in the pupal period and placed into single-sex vials to eclose. After eclosion, adult flies were maintained at a density of twenty flies per vial in single-sex groups. Unless otherwise stated, all experiments used 5- to 7-day-old virgin flies.

### Adult weight

Groups of 10 flies were placed in preweighed 1.5 ml tubes (Diamed Lab Supplies, DIATEC610-2550) and weighed on an analytical balance (Mettler-Toledo, ME104).

## Method details

### Capillary feeder (CAFE) assay

One biological replicate consisted of measurements from one group of 10 flies. Flies were placed into a specialized 15 ml conical vial with access to two capillary tubes (A-M Systems; 626000), as previously described^87^. Briefly, capillary tubes were filled with fly food media containing nutrients equivalent to the 1S diet (20.5 g/L sucrose, 70.9 g/L D-glucose, 45.3 g/L yeast extract (Sigma-Aldrich; Y1625) in water) or 0S diet (45.3 g/L yeast extract in water). To prevent evaporation, approximately 0.5 μl of mineral oil was added to the top of each capillary tube. The vials were placed into fitted holes in the lid of a large plastic container. To maintain high humidity throughout the experiment, a shallow layer of water was poured into the base of the container. The meniscus of the fly food media was marked before the start of the experiment and again after 24 hr. The distance between the marks was used to quantify the volume of fly food media that was consumed by the 10 flies (1 mm=0.15 μl). To account for sex differences in body size, the volume of fly food consumed was normalized to the weight of individual flies.

### Whole-body triglyceride measurements

One biological replicate consisted of five flies. Flies were collected in a 1.5 ml tube and homogenized in 200-350 μl of 0.1% Tween (Amresco, 0777-1L) in 1X phosphate-buffered saline (PBS; Sigma-Aldrich, P5493) using 50 μl of glass beads (Sigma-Aldrich, Z250473) agitated at 8 m/s for 5 s (OMNI International BeadRuptor 24). Triglyceride concentration was measured using either Sigma kit (G7793, T2449, F6428) or Stanbio kit (FT7610, BD386a/d, BD386b) according to the manufacturer’s instructions and as described previously^4,5^ with the following modifications. For Sigma kit, 30 μl of homogenate was transferred in duplicate to a 96-well plate and heat treated at 70°C for 10 min (Bio-Rad T100 Thermal Cycler). Standard curve was prepared using glycerol standard (Sigma, G7793) for concentrations of 0 mg/ml, 0.125 mg/ml, 0.25 mg/ml, 0.5 mg/ml, 1 mg/ml, and 2 mg/ml. After heat treatment, 30 μl of each concentration of standard solution was transferred to the same 96-well plate in duplicate. 30 μl of triglyceride reagent (Sigma, T2449) was added to 1 set of samples/standard solutions and 0.1% Tween in 1x PBS was added to the other set of samples/standard solutions. After 40 min incubation at 37°C, the 96-well plate was centrifuged at 3800 rpm for 3 min at 4°C (VWR® Benchtop General Purpose Centrifuges, 10830-744) to precipitate cellular debris. 7.5 μl of supernatant was transferred to a 384-well plate-reader plate and 25 μl of free glycerol reagent (Sigma, F6428) was added to each well. Plate was incubated at 37°C for 5 min, then absorbance was read at 540 nm (Thermo Scientific – Multiskan FC Microplate Photometer). For the Stanbio kit, 10 μl of either homogenate or triglyceride standard (FT7610) was added to 190 μl of activated triglyceride reagent (Enzymatic Triglyceride Reagent, BD386a/d; Triglyceride Activator, Cat. No. BD386b) in the 96-well plate. After 15 min incubation at room temperature, the absorbance was read at 540 nm. Triglyceride concentration was used to calculate percent body fat.

### Lipidomics and metabolomics

#### Sample preparation

Abdominal carcasses were isolated from 5-day-old male and female *w*^1118^ flies maintained on the 1S or 0S diet after eclosion. One biological replicate consisted of 30 abdominal carcasses dissected from 5- to 7-day-old adult males and females; a total of 9 biological replicates were collected for each sex and condition. After dissection, each carcass was snap-frozen on dry ice in a 2 mL screw-cap microcentrifuge tube (Thermo Fisher Scientific, Waltham, MA) and stored at –80°C until further processing. For a dual-phase extraction, samples were homogenized with a Mini-Beadbeater-24 (Biospec Products, Bartlesville, United States) at 3800 rpm. A total of 333 μl 90% MeOH was added and two 15 sec homogenization cycles were performed with a 1 min rest on dry ice in between. The homogenized solution was mixed with 1000 μl of methyl tert-butyl ether and shaken for 5 min. To induce the phase separation, 317 μl of water was added. After mixing, the solution rested at room temperature for 10 min and then the two-layer solution was centrifuged at 14,000 rpm for 15 min at 4°C. For LC-MS analysis, the clear upper and lower layers were separated into new vials, dried in a SpeedVac at 20°C, and reconstituted in 70 μl of acetonitrile and isopropanol (1:1, v/v) and 70 μl of acetonitrile and water (1:1, v/v) mixed solvent, respectively. A method blank sample was also prepared following the same protocol but without *Drosophila* abdomens.

#### LC-MS/MS experiment

For metabolomics, hydrophilic interaction chromatography (HILIC) separation was performed on a SeQuant ZIC-HILIC column (50 mm × 2.1 mm, 5 μm, 200 Å) with an elution time of 8 min. Mobile phase A was 5% acetonitrile in water with 10 mM ammonium acetate (Thermo Fisher Scientific) with pH = 9.8, adjusted by ammonium hydroxide (MilliporeSigma), and mobile phase B was 5% water in acetonitrile with no buffer. The LC elution gradient was optimized using the BAGO software^189^ as follows: 0 min, 10% A; 1 min, 10% A; 2 min, 20% A; 5 min, 20% A; 6 min, 45% A; 7 min, 95% A; 8 min, 95% A. For lipidomics, reversed phase (RP) separation was achieved on a Waters UPLC Acquity BEH C18 Column (1.0 mm ×100 mm, 1.7 µm, 130 Å, Milford, MA, USA) with an elution time of 25 min. Mobile phase A was acetonitrile and water (6:4, v/v) with 2 mM ammonium formate (pH = 4.8, adjusted by formic acid), and mobile phase B was isopropanol and acetonitrile (9:1, v/v). LC elution gradient was optimized using the BAGO software as follows: 0 min, 5% B; 1 min, 5% B; 4 min, 35% B; 7 min, 40% B; 10 min, 50% B; 13 min, 55% B; 16 min, 55% B; 19 min, 75% B; 22 min, 75% B; 25 min, 95% B.

The MS was operated in electrospray ionization (ESI) positive mode. The ESI source conditions were set as follows: dry gas temperature, 220°C; dry gas flow, 7 L/min; nebulizer gas pressure, 1.6 bar; capillary voltage, 4500 V for positive mode. The MS1 analysis was conducted using following parameters: mass range, 70-1000 *m/z*; spectrum type: centroid, calculated using maximum intensity; absolute intensity threshold: 250.

Data-dependent MS/MS analysis parameters: collision energy: 16-30 eV; cycle time: 3 s; spectra rate: 4 Hz when intensity < 10^4^ and 12 Hz when intensity > 10^5^, linearly increased from 10^4^ to 10^5^. External calibration was applied using sodium formate to ensure the *m/z* accuracy before sample analysis.

#### Data processing

The raw MS data were converted to ABF format in Reifycs Abf Converter (ver. 4.0.0). The converted data were processed in MS-DIAL (ver. 4.90) for feature extraction, feature alignment, and compound identification. The data processing parameters in MS-DIAL were set as follows: MS1 tolerance, 0.01 Da; MS/MS tolerance, 0.05 Da; mass slice width, 0.05 Da; smoothing method, linear weighted moving average; smoothing level, 3 scans; minimum peak width, 5 scans. Compound identification was achieved using NIST 20 for metabolomics and the internal library within MS-DIAL for lipidomics. Post-acquisition sample normalization was performed using the *MAFFIN* R package^210^ to ensure fair quantitative comparison.

#### Citrate assay

For females, one biological replicate consisted of ten fat bodies; for males one biological replicate consisted of fifteen fat bodies. Tissues were collected in a 1.5 ml tube and homogenized in 100 μl of 1XPBS (PBS; Sigma-Aldrich, P5493) using 50 μl of glass beads (Sigma-Aldrich, Z250473) agitated at 8 m/s for 5 s (OMNI International BeadRuptor 24).

Citrate concentration was measured using citrate assay kit (Sigma-Aldrich MAK333) according to the manufacturer’s instructions. Briefly, the homogenized solution was centrifuged at 14,000 rpm for 5 min at 4°C (VWR® Benchtop General Purpose Centrifuges, 10830-744) to precipitate cellular debris. 20 μl of supernatant was transferred in duplicate to a clear flat bottom 96-well plate, to which either 80 μl of reaction mix (85 μl of developer (#MAK333A) + 1 μl of CL enzyme solution (#MAK333B) + 1 μl of dye reagent (#MAK333C) + 1 μl of ODC enzyme (#MAK333D)) or blank reaction mix (85 μl of developer (#MAK333A) + 1 μl of dye reagent (#MAK333C) + 1 μl of ODC enzyme (#MAK333D)) were added. Standard curve was prepared using citrate standard (#MAK333E) for concentrations of 0 μM, 30 μM, 60 μM, 120 μM, 240 μM, and 400 μM. 20 μL of each standard solution was added to the 96 well plate, to which 80 μl of reaction mix was added. The whole plate was incubated at room temperature for 15 min and the absorbance was measured at 570 nm. The citrate levels were normalized to the protein concentration of each sample. Protein concentration was determined by Bradford assay (Bio-Rad #5000204), as we described previously^5^.

#### Excrement glucose measurements

One biological replicate consisted of 30 female flies or 60 male flies. Flies were fed with 1S diet containing 5% blue dye (FD&C Blue no.1, Ward’s Science, 470300-996) for 3.5 hr, and subsequently transferred into a 1.5 mL tube for excrement collection at room temperature. After 6 hr in the 1.5 mL tube, flies were discarded and 100 μL of 1x PBS was added to each tube; tubes were vortexed to dissolve and pool all the blue-stained excrement deposited in the tube. For blue dye measurement, 10 µL of the excrement was added to 10 µL of 1x PBS in 384-well plate and absorbance was measured at 630 nm (Thermo Scientific – Multiskan FC Microplate Photometer). The rest of the excreta (90 μL) was added to 110 µL of Glucose Reagent (Sigma-Aldrich, GAHK20) in 96-well plate and incubated overnight at 37 °C for measurement of glucose levels at 340 nm (Tecan Austria GmbH-Tecan Spark). Glucose content in the excrement was normalized with blue dye to account for sex differences in excreta volume, and to account for the number of flies that deposited the excreta^213^.

#### RNA extraction, cDNA synthesis, and Quantitative real-time PCR (qPCR)

One biological replicate consisted of 10 adult heads or 15 adult fat bodies. Samples were homogenized in 500 μl of Trizol (Thermo Fisher Scientific; 15596018) and precipitated using isopropanol and 75% ethanol to extract RNA. Pelleted RNA was resuspended using 20-30 μl of molecular biology grade water (Corning, 46-000-CV) and stored at −80°C until use. Each experiment contained 4-7 biological replicates per sex, per genotype, and per diet; each experiment was repeated twice.

For cDNA synthesis, an equal amount of RNA per reaction was DNase-treated and reverse transcribed according to manufacturer’s instructions using the QuantiTect Reverse Transcription Kit (Qiagen, 205314). Relative mRNA transcript levels were quantified using qPCR as described previously^5^. Data were normalized to the average fold change of *Actin5C* and β*-tubulin*.

#### Immunostaining

Brains were dissected in cold 1× PBS and fixed in fresh 4% paraformaldehyde at room temperature for 40 min. The brains were washed with 1× PBS three times and washed in PBST (PBS + 0.1% Triton X-100) for 10 min. The samples were blocked for 1 hr at 4°C in PBST + 3% goat normal serum (Sigma-Aldrich; S26). Samples were incubated with rabbit anti-Dilp3 (1:200)^208^ in blocking solution overnight at 4°C with gentle agitation. Samples were washed three times with PBST and then incubated with secondary antibody (goat anti-rabbit Alexa Fluor 488; 1:500) overnight at 4°C with gentle agitation. Samples were washed with PBST three times, with Hoescht added to the final PBST wash for 30 min (Sigma-Aldrich 33342, 1:1000). Brains were washed for 10 min in 1× PBS, mounted using SlowFade Diamond Antifade mountant (Invitrogen; S36972), and imaged using a Leica SP5 laser scanning confocal microscope. To quantify signal, we manually segmented IPC clusters and used Z-Project “sum slices” method, measured RawIntDen of IPC clusters and background after subtracting background RawIntDen. Signal was normalized to IPC size and to number of slices. To quantify anti-Dilp3 levels 16-21 brain were analyzed per sex and per condition.

#### IPC morphology

Brains were dissected in cold 1× PBS and fixed in fresh 4% paraformaldehyde at room temperature for 40 min. Brains were washed with 1× PBS for 5 min, to which Hoescht was added to a final dilution of 1:1000 for a 10 min incubation, followed by a 5 min wash in 1× PBS. Samples were mounted in a 100% glucose solution. Images were acquired using a Leica SP5 laser scanning confocal microscope; we dissected five brains per sex.

#### IPC calcium level measurements

To isolate the IPCs, individual flies were removed from food and anesthetized briefly with CO_2_. Brains were dissected in cold 1× PBS. Samples were fixed in 4% paraformaldehyde (Electron Microscopy Science 15710) for 40 min, followed by two washes in cold 1× PBS. Samples were incubated with Hoechst (Sigma-Aldrich, 33342) at a concentration of 1:1000 for 30 min and mounted in 75% (w/v) sucrose solution. Images were captured using a LSI3-Leica SP5 laser scanning confocal microscope and processed using Fiji (ImageJ;^192^). To quantify IPC activity we used IPC-specific GAL4 driver *dilp2-GAL4*^123^ to drive expression of both *UAS-mRFP* and *UAS-CaLexA* (*UAS-LexA-VP16-NFAT*)^95^. GFP intensity in each IPC cluster was quantified by measuring the sum of pixel intensity within the region of interest for both GFP and RFP. GFP was normalized to RFP that was co-expressed in the IPC to account for inter-individual and/or sex differences in GAL4-driven transgene expression (ImageJ;^209^).

#### IPC electrophysiology

Loose, whole-cell cell-attached patch clamp recordings were performed *in vivo* as previously described^214,215^. Adult flies 4-8 days old were food deprived on 1% agar for 24 hr. Flies were either kept on agar (starved), moved to 500 mM D-glucose for 30 min, or moved to whole food for 2 hr prior to mounting. Flies were briefly anesthetized with CO_2_ and mounted by the cervix into a custom chamber. Nail polish was applied around the eyes and over the labellum, adhering the mouthparts to the chamber to prevent excessive movement. After recovering in humidity chamber for 45 min, the cuticle and air sacs covering the *pars intercerebralis* were removed and the head and exposed brain were immersed in Adult Hemolymph-Like (AHL) solution (108 mM NaCl, 5 mM KCl, 4 mM NaHCO_3_, 1 mM NaH_2_PO_4_, 5 mM HEPES, 15 mM ribose, 2 mM Ca^2+^, 8.2 mM Mg^2+^, pH 7.5). IPCs were identified by expression of *UAS-mCD8::GFP* and glass electrodes (1.5-3.5 MΩ) filled with AHL were used for all recordings, with patch resistances ranging from 50-500 MΩ. Spikes were recorded in voltage-clamp mode in Multiclamp 700B at 20 kHz. Recordings were passed through a low-pass filter at 2kHz and high-pass filter of 5 Hz, and then band-pass filtered between 2 and 2000 Hz with a Bessel (8-pole) filter. Event detection was performed in Clampfit for spike analysis, with a threshold set between −6 to −12 pA. The number of spikes in a 30-60 second window for a given cell were counted and converted to spikes/sec. Data from 6-8 flies were collected per group, with recordings from 1-3 IPCs per animal.

#### Insulin sensitivity test

To measure insulin sensitivity, we adapted a published protocol^79^. Briefly, abdominal carcasses were isolated from 5-day-old adult flies; one biological replicate consisted of 15 abdominal carcasses. The carcasses were placed in a dish with ice-cold Schneider’s *Drosophila* solution (21720024, Thermo Scientific) for 30 min. After this initial incubation, Schneider’s media was replaced with 2 mL of fresh Schneider’s *Drosophila* solution or fresh Schneider’s *Drosophila* solution with recombinant human insulin (final concentration 1 μM; I2643, Sigma-Aldrich) and incubated at room temperature for 30 min. After stimulation, carcasses were collected in a tube, snap-frozen, and used for RNA isolation and qPCR analysis.

#### Lifespan

After eclosion, unmated male and female flies were transferred to new vials of fresh food every 2–3 days until no living flies remained in the vial. Deaths were recorded when the flies were transferred to fresh food.

#### Female fecundity

One 5-day-old virgin female was placed with a group of three 5-day-old virgin *CS* males for 1 hr. After a 1 hr mating period, the males were removed and the females were allowed to lay eggs. Females were transferred into new vials for egg-laying every 24 hr; the total number of pupae in each vial was counted several days later.

#### Male fertility

One singly-housed 5-day-old virgin male was placed with a group of three 5-day-old virgin *CS* females and allowed to interact for 1 hr. After the 1 hr mating period, the male was removed from the vial and the females were allowed to lay eggs. Females were transferred into new vials for egg-laying every 24 hr; the total number of pupae in each vial was counted several days later.

#### Quantification and statistical analysis

Statistical analyses and data presentation were completed using GraphPad Prism 10 (GraphPad Software, San Diego, CA, USA) (Supplemental Table S2). All data were tested for normality using the Shapiro-Wilk test. Normally-distributed data was subjected to parametric tests, including Student’s *t*-test, one-way ANOVA followed by Tukey’s HSD, and two-way ANOVA followed by Bonferroni post-hoc test as appropriate. For non-normally-distributed data we used Mann-Whitney test and Kruskal-Wallis test followed by Dunn’s test. For two-way ANOVA involving data that do not satisfy the normality assumption, aligned rank transformation was first applied using the art() function from the ARTool R package^216^. Then, ANOVA was performed on the transformed data with the base R anova() function. Finally, the art.con() function from the ARTool package was used to extract the main as well as the interaction effects. Default parameters were used in each step of the analysis. For all statistical analyses, differences were considered significant if *p*<0.05.

## Supplemental Figure Legends

**Figure S1.** Dietary sugar levels change body fat but not food consumption in adult flies. Related to Figure 1. (A) Food consumption per unit of body weight was significantly higher in 5-day-old adult *w*^1118^ females than males (*p*=0.0483; Student’s *t*-test); n=19 biological replicates. (B) Whole-body triglyceride levels were significantly lower in 5-day-old *CS* females maintained on the 0S diet compared with females kept on the 1S diet (*p*=0.0181); no significant change in body fat was observed in males (*p*=0.0582; sex:diet interaction *p*=0.7239). Two-way ANOVA followed by Bonferroni post-hoc test; n=8 biological replicates. (C) Food consumption was not significantly affected by diet in either 5-day-old adult *w^1118^*females or males (females: *p*>0.9999; males: *p*>0.9999; sex:diet interaction *p*=0.8450). Two-way ANOVA followed by Bonferroni post-hoc test; n=10-19 biological replicates. Note: food consumption data in 1S diet was replotted from panel S1A as food intake data on 1S and 0S were collected in the same series of experiments. (D) Whole-body triglyceride levels were significantly lower in 5-day-old *w^1118^* adult females and males maintained on 0S with L-glucose diet compared with sex-matched flies kept on the 1S diet (females: *p*<0.0001; males: *p*<0.0001); the magnitude of the decrease in body fat was greater in females than in males (sex:diet interaction *p*=0.0029). Two-way ANOVA followed by Bonferroni post-hoc test; n=8 biological replicates. Note: the 1S triglyceride data was replotted from Figure 1E as they were part of the same large experiment. All data plotted as mean ± SEM. ns indicates not significant with *p*>0.05; * *p*<0.05, \*\*\*\**p*<0.0001.

**Figure S2.** Sugar-induced changes in lipid species in the adult fat body. Related to Figure 2. (A) Specific fatty acids are enriched within triglyceride species that show a sex bias in abundance in 5-day-old adult *w^1118^* female (purple) and male (green) flies maintained on 1S (*p*<0.01; Student’s t-test). For example, all triglycerides carrying fatty acid C28:2 showed a female bias in abundance. (B) Citrate levels in the fat body of 5-day-old adult *w^1118^* males showed a trend toward lower levels of this metabolite compared with females (*p*=0.0630, Student’s *t*-test); n=6-7 biological replicates. Data plotted as mean ± SEM. (C) Number of lipid species within each subclass that show higher abundance in either 1S (purple) or 0S (pink) in 5-day-old adult *w^1118^* females (*p*<0.01; Student’s *t*-test). For example, 17/21 TG-O species are more abundant in 0S than in 1S, whereas 25/27 PE species are more abundant in 1S than 0S. (D) Specific fatty acids are enriched within triglyceride species that show sugar-dependent regulation in 5-day-old adult *w^1118^* females (*p*<0.01; Student’s *t*-test); 1S (purple), 0S (pink). For example, 5/5 triglycerides that contain C28:2 are higher in the 1S diet, whereas 22/29 C14:0 are higher in 0S. (E) Number of lipid species within each subclass that show higher abundance in either 1S (green) or 0S (blue) in 5-day-old adult *w^1118^* males (*p*<0.01; Student’s *t*-test). For example, 4/6 TG species are more abundant in 1S than in 0S. (F) Citrate levels in the fat body were significantly lower in 5-day-old *w^1118^* adult females and males maintained on 0S diet compared with sex-matched flies kept on the 1S diet (females: *p*=0.0193; males: *p*=0.0409; sex:diet interaction *p*=0.8806). Two-way ANOVA followed by Bonferroni post-hoc test; n=6-8 biological replicates. Data plotted as mean ± SEM. Note: the 1S citrate data was replotted from Figure S2B as they were part of the same large experiment. TG: triglyceride; TG-O: ether-linked TG; DG: diglyceride; PC: phosphatidylcholine; PC-O: ether-linked PC; PE: phosphatidylethanolamine; PG: phosphatidylglycerol; PI: phosphatidylinositol;. Acyl-Car: acyl-carnitine; Cer: Ceramide; ST: sterol; PS: phosphatidylserine; PE-O or PE-P: ether-linked PE; NAE: N-acetylethanolamine; SE: sterol ester; Hex-Cer: hexosyl-ceramide.

**Figure S3.** Sex-specific regulation of *dilp3* mRNA and protein. Related to Figure 3. (A-D) Representative images of maximum Z-projections from brains of 5-day-old *w^1118^* flies labelled with anti-Dilp3. Females maintained on 1S (A) and 0S (B); males maintained on 1S (C) and 0S (D). Scale bar represents 10 μm. (E) Quantification of anti-Dilp3 levels in brains 5-day-old *w^1118^* males and females maintained on either 1S or 0S. (F, G) Representative images of maximum Z-projections from brains of 5-day-old *dilp2-GAL4>UAS-CaLexA,UAS-mRFP* females (F) and males (G) maintained on 1S showing GFP levels as a readout for IPC Ca^2+^ activity. Scale bar represents 50 μm. (H) Quantification of GFP levels in brains of 5-day-old *dilp2-GAL4>UAS-CaLexA,UAS-mRFP* males and females maintained on 1S. Quantification of GFP levels due to IPC-specific expression of *UAS-CaLexA* in the IPCs, normalized to RFP levels. Normalized GFP intensity was not significantly different between females and males in 1S diet (*p*=0.4782; Mann-Whitney test); n=15-23 animals. (I) Changing the sexual identity of the IPC by expressing Tra^F^ (*dilp2-GAL4>UAS-tra^F^*) did not significantly alter mRNA levels of IPC-derived *dilp3* in head (*p*=0.7859 and *p*=0.0460, respectively). One-way ANOVA followed by Tukey HSD; n=4 biological replicates. (J) Changing the sexual identity of the APC by expressing Tra^F^ (*Akh-GAL4>UAS-tra*^F^) did not significantly alter mRNA levels of IPC-derived *dilp3* in heads (*p*=0.7143 and *p*=0.0009, respectively). One-way ANOVA followed by Tukey HSD; n=6 biological replicates. (K) Changing the sexual identity of neurons and neuropeptide-expressing cells using Tra^F^(*elav-GAL4>UAS-tra*^F^) significantly increased mRNA levels of IPC-derived *dilp3* in heads (*p*<0.0001 and *p*<0.0001, respectively). One-way ANOVA followed by Tukey HSD; n=6 biological replicates. Data plotted as mean ± SEM. ns indicates not significant with *p*>0.05, * *p*<0.05, ** *p*<0.01, *** *p*<0.001, **** *p*<0.0001.

**Figure S4.** Validating IPC ablation using *dilp2-GAL4*. Related to Figure 4. (A, B) Representative images of maximum Z-projections in 5-day-old flies where *dilp2-GAL4* was used to drive expression of membrane-bound GFP (*UAS-mCD8::GFP*) to visualize IPC morphology; female (A) and male (B) flies are shown. Scale bar represents 50 μm. (C, D) Representative images of maximum Z-projections in 5-day-old flies where *dilp2-GAL4* was used to drive expression of nuclear RFP (*UAS-nRFP*) to assess colocalization with anti-Dilp3; female (C, CD, CDD) and male (D, DD, DDD) flies. Scale bar represents 100 μm. (E, F) mRNA levels of IPC-derived *dilp2* in heads were significantly lower in female (E; *p*=0.0001, Student’s *t*-test) and showed a trend of lower *dilp2* in male (F; *p*=0.06, Mann-Whitney test) *dilp2-GAL4>UAS-rpr; tub-GAL80^ts^* flies with heat shock at 29°C compared to sex-matched flies reared in room temperature; n=4-6 biological replicates. (G-L) Representative images of maximum Z-projections in 5-day-old flies with (I-L) and without (G, H) *dilp2-GAL4*-mediated IPC ablation at the adult stage via heat shock-induced *rpr* overexpression. Brains were labelled with anti-Dilp3 to assess IPC ablation. As expected, IPC are present in animals without heat shock; IPC are fully (I, J) or partially (K, L) absent in animals with heat shock. Scale bar represents 100 μm. (M) Sex difference in body fat represented as a percent of the male-female difference in the control *dilp2-GAL4>+,tub-GAL80^ts^*genotype. (N) Sex difference in body fat represented as a percent of the male-female difference in the *dilp2-GAL4>rpr,tub-GAL80^ts^* genotype without heat shock. (O) *dilp2-GAL4>+ and +>UAS-dilp3* females maintained on 0S had significantly lower body fat than genotype-matched females kept on 1S, a sugar-dependent reduction in body fat that was blocked in females with IPC-specific overexpression of Dilp3 (*dilp2-GAL4>UAS-dilp3*; 1S to 0S *p*<0.0001 [GAL4], *p*<0.0001 [UAS] and *p*=0.5298 [Dilp3 overexpression]). Two-way ANOVA followed by Bonferroni post-hoc test; n=8 biological replicates. (P) *dilp2-GAL4>+ and +>UAS-dilp3* males maintained on 0S had significantly lower body fat than genotype-matched females kept on 1S, a sugar-dependent reduction in body fat that was blocked in males with IPC-specific overexpression of Dilp3 (*dilp2-GAL4>UAS-dilp3*; *p*=0.0029, *p*=0.0452 and *p*=0.1426, respectively). Two-way ANOVA followed by Bonferroni post-hoc test; n=8 biological replicates. (Q) Whole-body triglyceride levels were significantly lower in 5-day-old adult *w^1118^* females maintained on a diet supplemented with 0.5 mg/mL 20-HE compared with control females (p=0.0067, Student’s t-test); n=8 biological replicates. Data plotted as mean ± SEM. * *p*<0.05, ** p<0.01, *** *p*<0.001, **** *p*<0.0001.

## References

1. Kühnlein, R.P. (2012). Lipid droplet-based storage fat metabolism in Drosophila: Thematic Review Series: Lipid Droplet Synthesis and Metabolism: from Yeast to Man. J. Lipid Res. 53, 1430–1436. 10.1194/jlr.R024299.

2. Kühnlein, R.P. (2011). The contribution of the *Drosophila* model to lipid droplet research. Prog. Lipid Res. 50, 348–356. 10.1016/j.plipres.2011.04.001.

3. Gutierrez, E., Wiggins, D., Fielding, B., and Gould, A.P. (2007). Specialized hepatocyte-like cells regulate Drosophila lipid metabolism. Nature 445, 275–280. 10.1038/nature05382.

4. Sieber, M.H., and Thummel, C.S. (2009). The DHR96 Nuclear Receptor Controls Triacylglycerol Homeostasis in Drosophila. Cell Metab. 10, 481–490. 10.1016/j.cmet.2009.10.010.

5. Wat, L.W., Chao, C., Bartlett, R., Buchanan, J.L., Millington, J.W., Chih, H.J., Chowdhury, Z.S., Biswas, P., Huang, V., Shin, L.J., et al. (2020). A role for triglyceride lipase brummer in the regulation of sex differences in Drosophila fat storage and breakdown. PLOS Biol. 18, e3000595. 10.1371/journal.pbio.3000595.

6. Kis, V., Barti, B., Lippai, M., and Sass, M. (2015). Specialized Cortex Glial Cells Accumulate Lipid Droplets in Drosophila melanogaster. PloS One 10, e0131250. 10.1371/journal.pone.0131250.

7. Villanueva, J.E., Livelo, C., Trujillo, A.S., Chandran, S., Woodworth, B., Andrade, L., Le, H.D., Manor, U., Panda, S., and Melkani, G.C. (2019). Time-restricted feeding restores muscle function in Drosophila models of obesity and circadian-rhythm disruption. Nat. Commun. 10, 2700. 10.1038/s41467-019-10563-9.

8. Chao, C.F., Pesch, Y.-Y., Yu, H., Wang, C., Aristizabal, M.J., Huan, T., Tanentzapf, G., and Rideout, E.J. (2024). An important role for triglyceride in regulating spermatogenesis. eLife 12. 10.7554/eLife.87523.3.

9. Sieber, M.H., and Spradling, A.C. (2015). Steroid Signaling Establishes a Female Metabolic State and Regulates SREBP to Control Oocyte Lipid Accumulation. Curr. Biol. 25, 993–1004. 10.1016/j.cub.2015.02.019.

10. Wat, L.W., Chowdhury, Z.S., Millington, J.W., Biswas, P., and Rideout, E.J. (2021). Sex determination gene transformer regulates the male-female difference in Drosophila fat storage via the adipokinetic hormone pathway. eLife 10, e72350. 10.7554/eLife.72350.

11. Schwasinger-Schmidt, T.E., Kachman, S.D., and Harshman, L.G. (2012). Evolution of starvation resistance in Drosophila melanogaster: measurement of direct and correlated responses to artificial selection. J. Evol. Biol. 25, 378–387. 10.1111/j.1420-9101.2011.02428.x.

12. Buszczak, M., Lu, X., Segraves, W.A., Chang, T.Y., and Cooley, L. (2002). Mutations in the midway Gene Disrupt a Drosophila Acyl Coenzyme A: Diacylglycerol Acyltransferase. Genetics 160, 1511–1518. 10.1093/genetics/160.4.1511.

13. Wilfling, F., Wang, H., Haas, J.T., Krahmer, N., Gould, T.J., Uchida, A., Cheng, J.-X., Graham, M., Christiano, R., Fröhlich, F., et al. (2013). Triacylglycerol synthesis enzymes mediate lipid droplet growth by relocalizing from the ER to lipid droplets. Dev. Cell 24, 384–399. 10.1016/j.devcel.2013.01.013.

14. Bailey, A.P., Koster, G., Guillermier, C., Hirst, E.M.A., MacRae, J.I., Lechene, C.P., Postle, A.D., and Gould, A.P. (2015). Antioxidant Role for Lipid Droplets in a Stem Cell Niche of Drosophila. Cell 163, 340–353. 10.1016/j.cell.2015.09.020.

15. Ugrankar, R., Liu, Y., Provaznik, J., Schmitt, S., and Lehmann, M. (2011). Lipin is a central regulator of adipose tissue development and function in Drosophila melanogaster. Mol. Cell. Biol. 31, 1646–1656. 10.1128/MCB.01335-10.

16. Beller, M., Bulankina, A.V., Hsiao, H.-H., Urlaub, H., Jäckle, H., and Kühnlein, R.P. (2010). PERILIPIN-dependent control of lipid droplet structure and fat storage in Drosophila. Cell Metab. 12, 521–532. 10.1016/j.cmet.2010.10.001.

17. Grönke, S., Mildner, A., Fellert, S., Tennagels, N., Petry, S., Müller, G., Jäckle, H., and Kühnlein, R.P. (2005). Brummer lipase is an evolutionary conserved fat storage regulator in Drosophila. Cell Metab. 1, 323–330. 10.1016/j.cmet.2005.04.003.

18. Palm, W., Sampaio, J.L., Brankatschk, M., Carvalho, M., Mahmoud, A., Shevchenko, A., and Eaton, S. (2012). Lipoproteins in Drosophila melanogaster--assembly, function, and influence on tissue lipid composition. PLoS Genet. 8, e1002828. 10.1371/journal.pgen.1002828.

19. Bi, J., Xiang, Y., Chen, H., Liu, Z., Grönke, S., Kühnlein, R.P., and Huang, X. (2012). Opposite and redundant roles of the two Drosophila perilipins in lipid mobilization. J. Cell Sci. 125, 3568–3577. 10.1242/jcs.101329.

20. Tian, Y., Bi, J., Shui, G., Liu, Z., Xiang, Y., Liu, Y., Wenk, M.R., Yang, H., and Huang, X. (2011). Tissue-autonomous function of Drosophila seipin in preventing ectopic lipid droplet formation. PLoS Genet. 7, e1001364. 10.1371/journal.pgen.1001364.

21. Ding, L., Yang, X., Tian, H., Liang, J., Zhang, F., Wang, G., Wang, Y., Ding, M., Shui, G., and Huang, X. (2018). Seipin regulates lipid homeostasis by ensuring calcium-dependent mitochondrial metabolism. EMBO J. 37, e97572. 10.15252/embj.201797572.

22. Teixeira, L., Rabouille, C., Rørth, P., Ephrussi, A., and Vanzo, N.F. (2003). Drosophila Perilipin/ADRP homologue Lsd2 regulates lipid metabolism. Mech. Dev. 120, 1071–1081. 10.1016/s0925-4773(03)00158-8.

23. Grönke, S., Beller, M., Fellert, S., Ramakrishnan, H., Jäckle, H., and Kühnlein, R.P. (2003). Control of fat storage by a Drosophila PAT domain protein. Curr. Biol. CB 13, 603– 606. 10.1016/s0960-9822(03)00175-1.

24. Fauny, J.D., Silber, J., and Zider, A. (2005). Drosophila Lipid Storage Droplet 2 gene (Lsd-2) is expressed and controls lipid storage in wing imaginal discs. Dev. Dyn. Off. Publ. Am. Assoc. Anat. 232, 725–732. 10.1002/dvdy.20277.

25. Pospisilik, J.A., Schramek, D., Schnidar, H., Cronin, S.J.F., Nehme, N.T., Zhang, X., Knauf, C., Cani, P.D., Aumayr, K., Todoric, J., et al. (2010). Drosophila Genome-wide Obesity Screen Reveals Hedgehog as a Determinant of Brown versus White Adipose Cell Fate. Cell 140, 148–160. 10.1016/j.cell.2009.12.027.

26. Baumbach, J., Hummel, P., Bickmeyer, I., Kowalczyk, K.M., Frank, M., Knorr, K., Hildebrandt, A., Riedel, D., Jäckle, H., and Kühnlein, R.P. (2014). A Drosophila in vivo screen identifies store-operated calcium entry as a key regulator of adiposity. Cell Metab. 19, 331–343. 10.1016/j.cmet.2013.12.004.

27. Baranski, T.J., Kraja, A.T., Fink, J.L., Feitosa, M., Lenzini, P.A., Borecki, I.B., Liu, C.-T., Cupples, L.A., North, K.E., and Province, M.A. (2018). A high throughput, functional screen of human Body Mass Index GWAS loci using tissue-specific RNAi Drosophila melanogaster crosses. PLoS Genet. 14, e1007222. 10.1371/journal.pgen.1007222.

28. Kunte, A.S., Matthews, K.A., and Rawson, R.B. (2006). Fatty acid auxotrophy in Drosophila larvae lacking SREBP. Cell Metab. 3, 439–448. 10.1016/j.cmet.2006.04.011.

29. Teleman, A.A., Chen, Y.-W., and Cohen, S.M. (2005). 4E-BP functions as a metabolic brake used under stress conditions but not during normal growth. Genes Dev. 19, 1844– 1848. 10.1101/gad.341505.

30. Rajan, A., and Perrimon, N. (2012). Drosophila cytokine unpaired 2 regulates physiological homeostasis by remotely controlling insulin secretion. Cell 151, 123–137. 10.1016/j.cell.2012.08.019.

31. Reis, T., Gilst, M.R.V., and Hariharan, I.K. (2010). A Buoyancy-Based Screen of Drosophila Larvae for Fat-Storage Mutants Reveals a Role for Sir2 in Coupling Fat Storage to Nutrient Availability. PLOS Genet. 6, e1001206. 10.1371/journal.pgen.1001206.

32. Tsuda-Sakurai, K., Seong, K.-H., Horiuchi, J., Aigaki, T., and Tsuda, M. (2015). Identification of a novel role for Drosophila MESR4 in lipid metabolism. Genes Cells 20, 358–365. 10.1111/gtc.12221.

33. Bennick, R.A., Nagengast, A.A., and DiAngelo, J.R. (2019). The SR proteins SF2 and RBP1 regulate triglyceride storage in the fat body of *Drosophila*. Biochem. Biophys. Res. Commun. 516, 928–933. 10.1016/j.bbrc.2019.06.151.

34. Kishita, Y., Tsuda, M., and Aigaki, T. (2012). Impaired fatty acid oxidation in a Drosophila model of mitochondrial trifunctional protein (MTP) deficiency. Biochem. Biophys. Res. Commun. 419, 344–349. 10.1016/j.bbrc.2012.02.026.

35. Palanker, L., Tennessen, J.M., Lam, G., and Thummel, C.S. (2009). Drosophila HNF4 Regulates Lipid Mobilization and β-Oxidation. Cell Metab. 9, 228–239. 10.1016/j.cmet.2009.01.009.

36. Lee, S., Bao, H., Ishikawa, Z., Wang, W., and Lim, H.-Y. (2017). Cardiomyocyte Regulation of Systemic Lipid Metabolism by the Apolipoprotein B-Containing Lipoproteins in Drosophila. PLoS Genet. 13, e1006555. 10.1371/journal.pgen.1006555.

37. Parra-Peralbo, E., and Culi, J. (2011). Drosophila lipophorin receptors mediate the uptake of neutral lipids in oocytes and imaginal disc cells by an endocytosis-independent mechanism. PLoS Genet. 7, e1001297. 10.1371/journal.pgen.1001297.

38. Lim, H.-Y., Wang, W., Wessells, R.J., Ocorr, K., and Bodmer, R. (2011). Phospholipid homeostasis regulates lipid metabolism and cardiac function through SREBP signaling in Drosophila. Genes Dev. 25, 189–200. 10.1101/gad.1992411.

39. Bauer, R., Voelzmann, A., Breiden, B., Schepers, U., Farwanah, H., Hahn, I., Eckardt, F., Sandhoff, K., and Hoch, M. (2009). Schlank, a member of the ceramide synthase family controls growth and body fat in Drosophila. EMBO J. 28, 3706–3716. 10.1038/emboj.2009.305.

40. Li, Y., Hoffmann, J., Li, Y., Stephano, F., Bruchhaus, I., Fink, C., and Roeder, T. (2016). Octopamine controls starvation resistance, life span and metabolic traits in Drosophila. Sci. Rep. 6, 35359. 10.1038/srep35359.

41. Neckameyer, W.S., Coleman, C.M., Eadie, S., and Goodwin, S.F. (2007). Compartmentalization of neuronal and peripheral serotonin synthesis in Drosophila melanogaster. Genes Brain Behav. 6, 756–769. 10.1111/j.1601-183X.2007.00307.x.

42. Malita, A., Kubrak, O., Koyama, T., Ahrentløv, N., Texada, M.J., Nagy, S., Halberg, K.V., and Rewitz, K. (2022). A gut-derived hormone suppresses sugar appetite and regulates food choice in Drosophila. Nat. Metab. 4, 1532–1550. 10.1038/s42255-022-00672-z.

43. Yoshinari, Y., Kosakamoto, H., Kamiyama, T., Hoshino, R., Matsuoka, R., Kondo, S., Tanimoto, H., Nakamura, A., Obata, F., and Niwa, R. (2021). The sugar-responsive enteroendocrine neuropeptide F regulates lipid metabolism through glucagon-like and insulin-like hormones in Drosophila melanogaster. Nat. Commun. 12, 4818. 10.1038/s41467-021-25146-w.

44. Scopelliti, A., Bauer, C., Yu, Y., Zhang, T., Kruspig, B., Murphy, D.J., Vidal, M., Maddocks, O.D.K., and Cordero, J.B. (2019). A Neuronal Relay Mediates a Nutrient Responsive Gut/Fat Body Axis Regulating Energy Homeostasis in Adult Drosophila. Cell Metab. 29, 269–284.e10. 10.1016/j.cmet.2018.09.021.

45. Hentze, J.L., Carlsson, M.A., Kondo, S., Nässel, D.R., and Rewitz, K.F. (2015). The Neuropeptide Allatostatin A Regulates Metabolism and Feeding Decisions in Drosophila. Sci. Rep. 5, 11680. 10.1038/srep11680.

46. Kubrak, O., Koyama, T., Ahrentløv, N., Jensen, L., Malita, A., Naseem, M.T., Lassen, M., Nagy, S., Texada, M.J., Halberg, K.V., et al. (2022). The gut hormone Allatostatin C/Somatostatin regulates food intake and metabolic homeostasis under nutrient stress. Nat. Commun. 13, 692. 10.1038/s41467-022-28268-x.

47. Beshel, J., Dubnau, J., and Zhong, Y. (2017). A Leptin analog locally produced in the brain acts via a conserved neural circuit to modulate obesity-linked behaviors in Drosophila. Cell Metab. 25, 208–217. 10.1016/j.cmet.2016.12.013.

48. Yamamoto, R., Bai, H., Dolezal, A.G., Amdam, G., and Tatar, M. (2013). Juvenile hormone regulation of Drosophila aging. BMC Biol. 11, 85. 10.1186/1741-7007-11-85.

49. Kamoshida, Y., Fujiyama-Nakamura, S., Kimura, S., Suzuki, E., Lim, J., Shiozaki-Sato, Y., Kato, S., and Takeyama, K.-I. (2012). Ecdysone receptor (EcR) suppresses lipid accumulation in the Drosophila fat body via transcription control. Biochem. Biophys. Res. Commun. 421, 203–207. 10.1016/j.bbrc.2012.03.135.

50. Song, W., Veenstra, J.A., and Perrimon, N. (2014). Control of lipid metabolism by tachykinin in Drosophila. Cell Rep. 9, 40–47. 10.1016/j.celrep.2014.08.060.

51. Gera, J., Agard, M., Nave, H., Sajadi, F., Thorat, L., Kondo, S., Nässel, D.R., Paluzzi, J.-P.V., and Zandawala, M. (2024). Anti-diuretic hormone ITP signals via a guanylate cyclase receptor to modulate systemic homeostasis in Drosophila. eLife 13. 10.7554/eLife.97043.1.

52. Gáliková, M., and Klepsatel, P. (2022). Ion transport peptide regulates energy intake, expenditure, and metabolic homeostasis in Drosophila. Genetics 222, iyac150. 10.1093/genetics/iyac150.

53. DiAngelo, J.R., and Birnbaum, M.J. (2009). Regulation of Fat Cell Mass by Insulin in Drosophila melanogaster. Mol. Cell. Biol. 29, 6341–6352. 10.1128/MCB.00675-09.

54. Grönke, S., Clarke, D.-F., Broughton, S., Andrews, T.D., and Partridge, L. (2010). Molecular evolution and functional characterization of Drosophila insulin-like peptides. PLoS Genet. 6, e1000857. 10.1371/journal.pgen.1000857.

55. Grönke, S., Müller, G., Hirsch, J., Fellert, S., Andreou, A., Haase, T., Jäckle, H., and Kühnlein, R.P. (2007). Dual lipolytic control of body fat storage and mobilization in Drosophila. PLoS Biol. 5, e137. 10.1371/journal.pbio.0050137.

56. Bharucha, K.N., Tarr, P., and Zipursky, S.L. (2008). A glucagon-like endocrine pathway in Drosophila modulates both lipid and carbohydrate homeostasis. J. Exp. Biol. 211, 3103–3110. 10.1242/jeb.016451.

57. Gáliková, M., Diesner, M., Klepsatel, P., Hehlert, P., Xu, Y., Bickmeyer, I., Predel, R., and Kühnlein, R.P. (2015). Energy Homeostasis Control in Drosophila Adipokinetic Hormone Mutants. Genetics 201, 665–683. 10.1534/genetics.115.178897.

58. Zhao, X., and Karpac, J. (2017). Muscle Directs Diurnal Energy Homeostasis through a Myokine-Dependent Hormone Module in Drosophila. Curr. Biol. CB 27, 1941–1955.e6. 10.1016/j.cub.2017.06.004.

59. Kapan, N., Lushchak, O.V., Luo, J., and Nässel, D.R. (2012). Identified peptidergic neurons in the Drosophila brain regulate insulin-producing cells, stress responses and metabolism by coexpressed short neuropeptide F and corazonin. Cell. Mol. Life Sci. CMLS 69, 4051–4066. 10.1007/s00018-012-1097-z.

60. Luo, J., Becnel, J., Nichols, C.D., and Nässel, D.R. (2012). Insulin-producing cells in the brain of adult Drosophila are regulated by the serotonin 5-HT1A receptor. Cell. Mol. Life Sci. CMLS 69, 471–484. 10.1007/s00018-011-0789-0.

61. Zandawala, M., Yurgel, M.E., Texada, M.J., Liao, S., Rewitz, K.F., Keene, A.C., and Nässel, D.R. (2018). Modulation of Drosophila post-feeding physiology and behavior by the neuropeptide leucokinin. PLoS Genet. 14, e1007767. 10.1371/journal.pgen.1007767.

62. Kubrak, O., Jørgensen, A.F., Koyama, T., Lassen, M., Nagy, S., Hald, J., Mazzoni, G., Madsen, D., Hansen, J.B., Larsen, M.R., et al. (2024). LGR signaling mediates muscle-adipose tissue crosstalk and protects against diet-induced insulin resistance. Nat. Commun. 15, 6126. 10.1038/s41467-024-50468-w.

63. Heier, C., and Kühnlein, R.P. (2018). Triacylglycerol Metabolism in Drosophila melanogaster. Genetics 210, 1163–1184. 10.1534/genetics.118.301583.

64. Lee, G., and Park, J.H. (2004). Hemolymph sugar homeostasis and starvation-induced hyperactivity affected by genetic manipulations of the adipokinetic hormone-encoding gene in Drosophila melanogaster. Genetics 167, 311–323. 10.1534/genetics.167.1.311.

65. Noyes, B.E., Katz, F.N., and Schaffer, M.H. (1995). Identification and expression of the Drosophila adipokinetic hormone gene. Mol. Cell. Endocrinol. 109, 133–141. 10.1016/0303-7207(95)03492-p.

66. Mochanová, M., Tomčala, A., Svobodová, Z., and Kodrík, D. (2018). Role of adipokinetic hormone during starvation in Drosophila. Comp. Biochem. Physiol. B Biochem. Mol. Biol. 226, 26–35. 10.1016/j.cbpb.2018.08.004.

67. He, Q., Du, J., Wei, L., and Zhao, Z. (2020). AKH-FOXO pathway regulates starvation-induced sleep loss through remodeling of the small ventral lateral neuron dorsal projections. PLoS Genet. 16, e1009181. 10.1371/journal.pgen.1009181.

68. Li, H., Janssens, J., De Waegeneer, M., Kolluru, S.S., Davie, K., Gardeux, V., Saelens, W., David, F., Brbić, M., Spanier, K., et al. (2022). Fly Cell Atlas: a single-nucleus transcriptomic atlas of the adult fruit fly. Science 375, eabk2432. 10.1126/science.abk2432.

69. Leader, D.P., Krause, S.A., Pandit, A., Davies, S.A., and Dow, J.A.T. (2018). FlyAtlas 2: a new version of the Drosophila melanogaster expression atlas with RNA-Seq, miRNA-Seq and sex-specific data. Nucleic Acids Res. 46, D809–D815. 10.1093/nar/gkx976.

70. De Groef, S., Ribeiro Lopes, M., Winant, M., Rosschaert, E., Wilms, T., Bolckmans, L., Calevro, F., and Callaerts, P. (2024). Reference genes to study the sex-biased expression of genes regulating Drosophila metabolism. Sci. Rep. 14, 9518. 10.1038/s41598-024-58863-5.

71. Ja, W.W., Carvalho, G.B., Mak, E.M., de la Rosa, N.N., Fang, A.Y., Liong, J.C., Brummel, T., and Benzer, S. (2007). Prandiology of Drosophila and the CAFE assay. Proc. Natl. Acad. Sci. U. S. A. 104, 8253–8256. 10.1073/pnas.0702726104.

72. Al-Anzi, B., Sapin, V., Waters, C., Zinn, K., Wyman, R.J., and Benzer, S. (2009). Obesity-Blocking Neurons in Drosophila. Neuron 63, 329–341. 10.1016/j.neuron.2009.07.021.

73. Tennessen, J.M., Barry, W.E., Cox, J., and Thummel, C.S. (2014). Methods for studying metabolism in Drosophila. Methods San Diego Calif 68, 105–115. 10.1016/j.ymeth.2014.02.034.

74. Hildebrandt, A., Bickmeyer, I., and Kühnlein, R.P. (2011). Reliable Drosophila body fat quantification by a coupled colorimetric assay. PloS One 6, e23796. 10.1371/journal.pone.0023796.

75. De Groef, S., Wilms, T., Balmand, S., Calevro, F., and Callaerts, P. (2021). Sexual Dimorphism in Metabolic Responses to Western Diet in Drosophila melanogaster. Biomolecules 12, 33. 10.3390/biom12010033.

76. Musselman, L.P., Fink, J.L., and Baranski, T.J. (2019). Similar effects of high-fructose and high-glucose feeding in a Drosophila model of obesity and diabetes. PLoS ONE 14, e0217096. 10.1371/journal.pone.0217096.

77. Rovenko, B.M., Perkhulyn, N.V., Gospodaryov, D.V., Sanz, A., Lushchak, O.V., and Lushchak, V.I. (2015). High consumption of fructose rather than glucose promotes a diet-induced obese phenotype in *Drosophila melanogaster*. Comp. Biochem. Physiol. A. Mol. Integr. Physiol. 180, 75–85. 10.1016/j.cbpa.2014.11.008.

78. van Dam, E., van Leeuwen, L.A.G., dos Santos, E., James, J., Best, L., Lennicke, C., Vincent, A.J., Marinos, G., Foley, A., Buricova, M., et al. (2020). Sugar-Induced Obesity and Insulin Resistance Are Uncoupled from Shortened Survival in Drosophila. Cell Metab. 31, 710–725.e7. 10.1016/j.cmet.2020.02.016.

79. Musselman, L.P., Fink, J.L., Narzinski, K., Ramachandran, P.V., Sukumar Hathiramani, S., Cagan, R.L., and Baranski, T.J. (2011). A high-sugar diet produces obesity and insulin resistance in wild-type Drosophila. Dis. Model. Mech. 4, 842–849. 10.1242/dmm.007948.

80. Millington, J.W., Biswas, P., Chao, C., Xia, Y.H., Wat, L.W., Brownrigg, G.P., Sun, Z., Basner-Collins, P.J., Klein Geltink, R.I., and Rideout, E.J. (2022). A low-sugar diet enhances Drosophila body size in males and females via sex-specific mechanisms. Dev. Camb. Engl. 149, dev200491. 10.1242/dev.200491.

81. Lewis, E.B. (1960). A new standard food medium. Drosoph. Inf. Serv. 34, 1–55.

82. Nayak, N., and Mishra, M. (2021). High fat diet induced abnormalities in metabolism, growth, behavior, and circadian clock in *Drosophila melanogaster*. Life Sci. 281, 119758. 10.1016/j.lfs.2021.119758.

83. Birse, R.T., Choi, J., Reardon, K., Rodriguez, J., Graham, S., Diop, S., Ocorr, K., Bodmer, R., and Oldham, S. (2010). High-fat-diet-induced obesity and heart dysfunction are regulated by the TOR pathway in Drosophila. Cell Metab. 12, 533–544. 10.1016/j.cmet.2010.09.014.

84. Heinrichsen, E.T., and Haddad, G.G. (2012). Role of High-Fat Diet in Stress Response of Drosophila. PLOS ONE 7, e42587. 10.1371/journal.pone.0042587.

85. Woodcock, K.J., Kierdorf, K., Pouchelon, C.A., Vivancos, V., Dionne, M.S., and Geissmann, F. (2015). Macrophage-Derived upd3 Cytokine Causes Impaired Glucose Homeostasis and Reduced Lifespan in Drosophila Fed a Lipid-Rich Diet. Immunity 42, 133–144. 10.1016/j.immuni.2014.12.023.

86. Liao, S., Amcoff, M., and Nässel, D.R. (2021). Impact of high-fat diet on lifespan, metabolism, fecundity and behavioral senescence in *Drosophila*. Insect Biochem. Mol. Biol. 133, 103495. 10.1016/j.ibmb.2020.103495.

87. Stafford, J.W., Lynd, K.M., Jung, A.Y., and Gordon, M.D. (2012). Integration of Taste and Calorie Sensing in Drosophila. J. Neurosci. 32, 14767–14774. 10.1523/JNEUROSCI.1887-12.2012.

88. Regan, J.C., Khericha, M., Dobson, A.J., Bolukbasi, E., Rattanavirotkul, N., and Partridge, L. (2016). Sex difference in pathology of the ageing gut mediates the greater response of female lifespan to dietary restriction. eLife 5, e10956. 10.7554/eLife.10956.

89. Mirth, C.K., and Riddiford, L.M. (2007). Size assessment and growth control: how adult size is determined in insects. BioEssays News Rev. Mol. Cell. Dev. Biol. 29, 344–355. 10.1002/bies.20552.

90. Kréneisz, O., Chen, X., Fridell, Y.-W.C., and Mulkey, D.K. (2010). Glucose increases activity and Ca2+ in insulin-producing cells of adult Drosophila. Neuroreport 21, 1116– 1120. 10.1097/WNR.0b013e3283409200.

91. Nässel, D.R., Liu, Y., and Luo, J. (2015). Insulin/IGF signaling and its regulation in *Drosophila*. Gen. Comp. Endocrinol. 221, 255–266. 10.1016/j.ygcen.2014.11.021.

92. Park, S., Alfa, R.W., Topper, S.M., Kim, G.E.S., Kockel, L., and Kim, S.K. (2014). A Genetic Strategy to Measure Circulating Drosophila Insulin Reveals Genes Regulating Insulin Production and Secretion. PLOS Genet. 10, e1004555. 10.1371/journal.pgen.1004555.

93. Oh, Y., Lai, J.S.-Y., Mills, H.J., Erdjument-Bromage, H., Giammarinaro, B., Saadipour, K., Wang, J.G., Abu, F., Neubert, T.A., and Suh, G.S.B. (2019). A glucose-sensing neuron pair regulates insulin and glucagon in Drosophila. Nature 574, 559–564. 10.1038/s41586-019-1675-4.

94. Buhler, K., Clements, J., Winant, M., Bolckmans, L., Vulsteke, V., and Callaerts, P. (2018). Growth control through regulation of insulin signalling by nutrition-activated steroid hormone in Drosophila. Development 145, dev165654. 10.1242/dev.165654.

95. Masuyama, K., Zhang, Y., Rao, Y., and Wang, J.W. (2012). Mapping Neural Circuits with Activity-Dependent Nuclear Import of a Transcription Factor. J. Neurogenet. 26, 89–102. 10.3109/01677063.2011.642910.

96. Bisen, R.S., Iqbal, F.M., Cascino-Milani, F., Bockemühl, T., and Ache, J.M. (2024). Nutritional state-dependent modulation of Insulin-Producing Cells in Drosophila. eLife 13. 10.7554/eLife.98514.1.

97. Sturtevant, A.H. (1945). A Gene in Drosophila Melanogaster That Transforms Females into Males. Genetics 30, 297–299. 10.1093/genetics/30.3.297.

98. McKeown, M., Belote, J.M., and Baker, B.S. (1987). A molecular analysis of *transformer*, a gene in drosophila melanogaster that controls female sexual differentiation. Cell 48, 489–499. 10.1016/0092-8674(87)90199-1.

99. Camara, N., Whitworth, C., and Van Doren, M. (2008). The creation of sexual dimorphism in the Drosophila soma. Curr. Top. Dev. Biol. 83, 65–107. 10.1016/S0070-2153(08)00403-1.

100. Oliver, B. (2002). Genetic control of germline sexual dimorphism in Drosophila. In International Review of Cytology, K. W. Jeon, ed. (Academic Press), pp. 1–60. 10.1016/S0074-7696(02)19010-3.

101. Burtis, K.C. (1993). The regulation of sex determination and sexually dimorphic differentiation in Drosophila. Curr. Opin. Cell Biol. 5, 1006–1014. 10.1016/0955-0674(93)90085-5.

102. Rideout, E.J., Dornan, A.J., Neville, M.C., Eadie, S., and Goodwin, S.F. (2010). Control of sexual differentiation and behavior by the doublesex gene in Drosophila melanogaster. Nat. Neurosci. 13, 458–466. 10.1038/nn.2515.

103. Rideout, E.J., Narsaiya, M.S., and Grewal, S.S. (2015). The Sex Determination Gene transformer Regulates Male-Female Differences in Drosophila Body Size. PLOS Genet. 11, e1005683. 10.1371/journal.pgen.1005683.

104. Hudry, B., Khadayate, S., and Miguel-Aliaga, I. (2016). The sexual identity of adult intestinal stem cells controls organ size and plasticity. Nature 530, 344–348. 10.1038/nature16953.

105. Hudry, B., de Goeij, E., Mineo, A., Gaspar, P., Hadjieconomou, D., Studd, C., Mokochinski, J.B., Kramer, H.B., Plaçais, P.-Y., Preat, T., et al. (2019). Sex Differences in Intestinal Carbohydrate Metabolism Promote Food Intake and Sperm Maturation. Cell 178, 901–918.e16. 10.1016/j.cell.2019.07.029.

106. Evans, D.S., and Cline, T.W. (2007). Drosophila melanogaster Male Somatic Cells Feminized Solely by TraF Can Collaborate With Female Germ Cells to Make Functional Eggs. Genetics 175, 631–642. 10.1534/genetics.106.066332.

107. Regan, J.C., Lu, Y.-X., Ureña, E., Meilenbrock, R.L., Catterson, J.H., Kißler, D., Fröhlich, J., Funk, E., and Partridge, L. (2022). Sexual identity of enterocytes regulates autophagy to determine intestinal health, lifespan and responses to rapamycin. Nat. Aging 2, 1145–1158. 10.1038/s43587-022-00308-7.

108. Kim, J., and Neufeld, T.P. (2015). Dietary sugar promotes systemic TOR activation in Drosophila through AKH-dependent selective secretion of Dilp3. Nat. Commun. 6, 6846. 10.1038/ncomms7846.

109. Birse, R.T., Söderberg, J.A.E., Luo, J., Winther, Å.M.E., and Nässel, D.R. (2011). Regulation of insulin-producing cells in the adult Drosophila brain via the tachykinin peptide receptor DTKR. J. Exp. Biol. 214, 4201–4208. 10.1242/jeb.062091.

110. Brogiolo, W., Stocker, H., Ikeya, T., Rintelen, F., Fernandez, R., and Hafen, E. (2001). An evolutionarily conserved function of the *Drosophila* insulin receptor and insulin-like peptides in growth control. Curr. Biol. 11, 213–221. 10.1016/S0960-9822(01)00068-9.

111. Ikeya, T., Galic, M., Belawat, P., Nairz, K., and Hafen, E. (2002). Nutrient-dependent expression of insulin-like peptides from neuroendocrine cells in the CNS contributes to growth regulation in Drosophila. Curr. Biol. CB 12, 1293–1300. 10.1016/s0960-9822(02)01043-6.

112. Alic, N., Andrews, T.D., Giannakou, M.E., Papatheodorou, I., Slack, C., Hoddinott, M.P., Cochemé, H.M., Schuster, E.F., Thornton, J.M., and Partridge, L. (2011). Genome-wide dFOXO targets and topology of the transcriptomic response to stress and insulin signalling. Mol. Syst. Biol. 7, 502. 10.1038/msb.2011.36.

113. Puig, O., and Tjian, R. (2005). Transcriptional feedback control of insulin receptor by dFOXO/FOXO1. Genes Dev. 19, 2435–2446. 10.1101/gad.1340505.

114. Puig, O., Marr, M.T., Ruhf, M.L., and Tjian, R. (2003). Control of cell number by Drosophila FOXO: downstream and feedback regulation of the insulin receptor pathway. Genes Dev. 17, 2006–2020. 10.1101/gad.1098703.

115. Zinke, I., Schütz, C.S., Katzenberger, J.D., Bauer, M., and Pankratz, M.J. (2002). Nutrient control of gene expression in Drosophila: microarray analysis of starvation and sugar-dependent response. EMBO J. 21, 6162–6173. 10.1093/emboj/cdf600.

116. Macotela, Y., Boucher, J., Tran, T.T., and Kahn, C.R. (2009). Sex and depot differences in adipocyte insulin sensitivity and glucose metabolism. Diabetes 58, 803–812. 10.2337/db08-1054.

117. Geer, E.B., and Shen, W. (2009). Gender differences in insulin resistance, body composition, and energy balance. Gend. Med. 6 *Suppl 1*, 60–75. 10.1016/j.genm.2009.02.002.

118. Engel, F.L., and White, J.E. (1960). Some Hormonal Influences on Fat Mobilization from Adipose Tissue. Am. J. Clin. Nutr. 8, 691–704. 10.1093/ajcn/8.5.691.

119. Zhao, J., Wu, Y., Rong, X., Zheng, C., and Guo, J. (2020). Anti-Lipolysis Induced by Insulin in Diverse Pathophysiologic Conditions of Adipose Tissue. Diabetes Metab. Syndr. Obes. Targets Ther. 13, 1575–1585. 10.2147/DMSO.S250699.

120. Chakrabarti, P., Kim, J.Y., Singh, M., Shin, Y.-K., Kim, J., Kumbrink, J., Wu, Y., Lee, M.-J., Kirsch, K.H., Fried, S.K., et al. (2013). Insulin Inhibits Lipolysis in Adipocytes via the Evolutionarily Conserved mTORC1-Egr1-ATGL-Mediated Pathway. Mol. Cell. Biol. 33, 3659–3666. 10.1128/MCB.01584-12.

121. Scherer, T., O’Hare, J., Diggs-Andrews, K., Schweiger, M., Cheng, B., Lindtner, C., Zielinski, E., Vempati, P., Su, K., Dighe, S., et al. (2011). Brain Insulin Controls Adipose Tissue Lipolysis and Lipogenesis. Cell Metab. 13, 183–194. 10.1016/j.cmet.2011.01.008.

122. Krycer, J.R., Quek, L.-E., Francis, D., Zadoorian, A., Weiss, F.C., Cooke, K.C., Nelson, M.E., Diaz-Vegas, A., Humphrey, S.J., Scalzo, R., et al. (2020). Insulin signaling requires glucose to promote lipid anabolism in adipocytes. J. Biol. Chem. 295, 13250– 13266. 10.1074/jbc.RA120.014907.

123. Rulifson, E.J., Kim, S.K., and Nusse, R. (2002). Ablation of Insulin-Producing Neurons in Flies: Growth and Diabetic Phenotypes. Science 296, 1118–1120. 10.1126/science.1070058.

124. Clancy, D.J., Gems, D., Harshman, L.G., Oldham, S., Stocker, H., Hafen, E., Leevers, S.J., and Partridge, L. (2001). Extension of life-span by loss of CHICO, a Drosophila insulin receptor substrate protein. Science 292, 104–106. 10.1126/science.1057991.

125. Millington, J.W., Brownrigg, G.P., Basner-Collins, P.J., Sun, Z., and Rideout, E.J. (2021). Genetic manipulation of insulin/insulin-like growth factor signaling pathway activity has sex-biased effects on Drosophila body size. G3 GenesGenomesGenetics 11, jkaa067. 10.1093/g3journal/jkaa067.

126. Grewal, S.S. (2009). Insulin/TOR signaling in growth and homeostasis: a view from the fly world. Int. J. Biochem. Cell Biol. 41, 1006–1010. 10.1016/j.biocel.2008.10.010.

127. Teleman, A.A. (2009). Molecular mechanisms of metabolic regulation by insulin in Drosophila. Biochem. J. 425, 13–26. 10.1042/BJ20091181.

128. Osterwalder, T., Yoon, K.S., White, B.H., and Keshishian, H. (2001). A conditional tissue-specific transgene expression system using inducible GAL4. Proc. Natl. Acad. Sci. U. S. A. 98, 12596–12601. 10.1073/pnas.221303298.

129. Roman, G., Endo, K., Zong, L., and Davis, R.L. (2001). P{Switch}, a system for spatial and temporal control of gene expression in Drosophila melanogaster. Proc. Natl. Acad. Sci. 98, 12602–12607. 10.1073/pnas.221303998.

130. Poirier, L., Shane, A., Zheng, J., and Seroude, L. (2008). Characterization of the Drosophila Gene-Switch system in aging studies: a cautionary tale. Aging Cell 7, 758–770. 10.1111/j.1474-9726.2008.00421.x.

131. Yamada, R., Deshpande, S.A., Keebaugh, E.S., Ehrlich, M.R., Soto Obando, A., and Ja, W.W. (2017). Mifepristone Reduces Food Palatability and Affects Drosophila Feeding and Lifespan. J. Gerontol. A. Biol. Sci. Med. Sci. 72, 173–180. 10.1093/gerona/glw072.

132. Landis, G.N., Salomon, M.P., Keroles, D., Brookes, N., Sekimura, T., and Tower, J. (2015). The progesterone antagonist mifepristone/RU486 blocks the negative effect on life span caused by mating in female Drosophila. Aging 7, 53–69. 10.18632/aging.100721.

133. McClure, C.D., Hassan, A., Aughey, G.N., Butt, K., Estacio-Gómez, A., Duggal, A., Ying Sia, C., Barber, A.F., and Southall, T.D. (2022). An auxin-inducible, GAL4-compatible, gene expression system for Drosophila. eLife 11, e67598. 10.7554/eLife.67598.

134. Fleck, S.A., Biswas, P., DeWitt, E.D., Knuteson, R.L., Eisman, R.C., Nemkov, T., D’Alessandro, A., Tennessen, J.M., Rideout, E., and Weaver, L.N. (2024). Auxin exposure disrupts feeding behavior and fatty acid metabolism in adult Drosophila. eLife 12, RP91953. 10.7554/eLife.91953.

135. McGuire, S.E., Le, P.T., Osborn, A.J., Matsumoto, K., and Davis, R.L. (2003). Spatiotemporal rescue of memory dysfunction in Drosophila. Science 302, 1765–1768. 10.1126/science.1089035.

136. Klepsatel, P., Gáliková, M., Xu, Y., and Kühnlein, R.P. (2016). Thermal stress depletes energy reserves in Drosophila. Sci. Rep. 6, 33667. 10.1038/srep33667.

137. Klepsatel, P., Wildridge, D., and Gáliková, M. (2019). Temperature induces changes in Drosophila energy stores. Sci. Rep. 9, 5239. 10.1038/s41598-019-41754-5.

138. Tanabe, K., Itoh, M., and Tonoki, A. (2017). Age-Related Changes in Insulin-like Signaling Lead to Intermediate-Term Memory Impairment in *Drosophila*. Cell Rep. 18, 1598–1605. 10.1016/j.celrep.2017.01.053.

139. Enell, L.E., Kapan, N., Söderberg, J.A.E., Kahsai, L., and Nässel, D.R. (2010). Insulin Signaling, Lifespan and Stress Resistance Are Modulated by Metabotropic GABA Receptors on Insulin Producing Cells in the Brain of Drosophila. PLOS ONE 5, e15780. 10.1371/journal.pone.0015780.

140. Corl, A.B., Rodan, A.R., and Heberlein, U. (2005). Insulin signaling in the nervous system regulates ethanol intoxication in Drosophila melanogaster. Nat. Neurosci. 8, 18–19. 10.1038/nn1363.

141. Haselton, A., Sharmin, E., Schrader, J., Sah, M., Poon, P., and Fridell, Y.-W.C. (2010). Partial ablation of adult Drosophila insulin-producing neurons modulates glucose homeostasis and extends life span without insulin resistance. Cell Cycle 9, 3063–3071. 10.4161/cc.9.15.12458.

142. Broughton, S.J., Piper, M.D.W., Ikeya, T., Bass, T.M., Jacobson, J., Driege, Y., Martinez, P., Hafen, E., Withers, D.J., Leevers, S.J., et al. (2005). Longer lifespan, altered metabolism, and stress resistance in Drosophila from ablation of cells making insulin-like ligands. Proc. Natl. Acad. Sci. U. S. A. 102, 3105–3110. 10.1073/pnas.0405775102.

143. Nässel, D.R., Kubrak, O.I., Liu, Y., Luo, J., and Lushchak, O.V. (2013). Factors that regulate insulin producing cells and their output in Drosophila. Front. Physiol. 4, 252. 10.3389/fphys.2013.00252.

144. Kimura, K.D., Tissenbaum, H.A., Liu, Y., and Ruvkun, G. (1997). daf-2, an insulin receptor-like gene that regulates longevity and diapause in Caenorhabditis elegans. Science 277, 942–946. 10.1126/science.277.5328.942.

145. Kenyon, C., Chang, J., Gensch, E., Rudner, A., and Tabtiang, R. (1993). A C. elegans mutant that lives twice as long as wild type. Nature 366, 461–464. 10.1038/366461a0.

146. Partridge, L., Alic, N., Bjedov, I., and Piper, M.D.W. (2011). Ageing in Drosophila: The role of the insulin/Igf and TOR signalling network. Exp. Gerontol. 46, 376–381. 10.1016/j.exger.2010.09.003.

147. Blüher, M., Kahn, B.B., and Kahn, C.R. (2003). Extended longevity in mice lacking the insulin receptor in adipose tissue. Science 299, 572–574. 10.1126/science.1078223.

148. Fontana, L., Partridge, L., and Longo, V.D. (2010). Extending healthy life span--from yeast to humans. Science 328, 321–326. 10.1126/science.1172539.

149. Kuo, T.-H., Fedina, T.Y., Hansen, I., Dreisewerd, K., Dierick, H.A., Yew, J.Y., and Pletcher, S.D. (2012). Insulin Signaling Mediates Sexual Attractiveness in Drosophila. PLoS Genet. 8, e1002684. 10.1371/journal.pgen.1002684.

150. Carvalho, G.B., Kapahi, P., Anderson, D.J., and Benzer, S. (2006). Allocrine Modulation of Feeding Behavior by the Sex Peptide of *Drosophila*. Curr. Biol. 16, 692–696. 10.1016/j.cub.2006.02.064.

151. Hadjieconomou, D., King, G., Gaspar, P., Mineo, A., Blackie, L., Ameku, T., Studd, C., de Mendoza, A., Diao, F., White, B.H., et al. (2020). Enteric neurons increase maternal food intake during reproduction. Nature 587, 455–459. 10.1038/s41586-020-2866-8.

152. Cognigni, P., Bailey, A.P., and Miguel-Aliaga, I. (2011). Enteric Neurons and Systemic Signals Couple Nutritional and Reproductive Status with Intestinal Homeostasis. Cell Metab. 13, 92–104. 10.1016/j.cmet.2010.12.010.

153. Ja, W.W., Carvalho, G.B., Zid, B.M., Mak, E.M., Brummel, T., and Benzer, S. (2009). Water- and nutrient-dependent effects of dietary restriction on Drosophila lifespan. Proc. Natl. Acad. Sci. U. S. A. 106, 18633–18637. 10.1073/pnas.0908016106.

154. Lee, K.P., Simpson, S.J., Clissold, F.J., Brooks, R., Ballard, J.W.O., Taylor, P.W., Soran, N., and Raubenheimer, D. (2008). Lifespan and reproduction in Drosophila: New insights from nutritional geometry. Proc. Natl. Acad. Sci. U. S. A. 105, 2498–2503. 10.1073/pnas.0710787105.

155. Chapman, T., and Partridge, L. (1996). Female fitness in Drosophila melanogaster: an interaction between the effect of nutrition and of encounter rate with males. Proc. Biol. Sci. 263, 755–759. 10.1098/rspb.1996.0113.

156. Barnes, A.I., Wigby, S., Boone, J.M., Partridge, L., and Chapman, T. (2008). Feeding, fecundity and lifespan in female Drosophila melanogaster. Proc. Biol. Sci. 275, 1675– 1683. 10.1098/rspb.2008.0139.

157. Grandison, R.C., Piper, M.D.W., and Partridge, L. (2009). Amino-acid imbalance explains extension of lifespan by dietary restriction in Drosophila. Nature 462, 1061–1064. 10.1038/nature08619.

158. Laturney, M., Sterne, G.R., and Scott, K. (2023). Mating activates neuroendocrine pathways signaling hunger in Drosophila females. eLife 12, e85117. 10.7554/eLife.85117.

159. Strilbytska, O., Yurkevych, I., Semaniuk, U., Gospodaryov, D., Simpson, S.J., and Lushchak, O. (2024). Life-History Trade-Offs in Drosophila: Flies Select a Diet to Maximize Reproduction at the Expense of Lifespan. J. Gerontol. Ser. A 79, glae057. 10.1093/gerona/glae057.

160. Lee, K.P., Kim, J.-S., and Min, K.-J. (2013). Sexual dimorphism in nutrient intake and life span is mediated by mating in *Drosophila melanogaster*. Anim. Behav. 86, 987–992. 10.1016/j.anbehav.2013.08.018.

161. Drummond-Barbosa, D., and Spradling, A.C. (2001). Stem cells and their progeny respond to nutritional changes during Drosophila oogenesis. Dev. Biol. 231, 265–278. 10.1006/dbio.2000.0135.

162. Camus, M.F., Huang, C.-C., Reuter, M., and Fowler, K. (2018). Dietary choices are influenced by genotype, mating status, and sex in Drosophila melanogaster. Ecol. Evol. 8, 5385–5393. 10.1002/ece3.4055.

163. Ribeiro, C., and Dickson, B.J. (2010). Sex peptide receptor and neuronal TOR/S6K signaling modulate nutrient balancing in Drosophila. Curr. Biol. CB 20, 1000–1005. 10.1016/j.cub.2010.03.061.

164. Hsu, H.-J., and Drummond-Barbosa, D. (2009). Insulin levels control female germline stem cell maintenance via the niche in Drosophila. Proc. Natl. Acad. Sci. 106, 1117–1121. 10.1073/pnas.0809144106.

165. Burn, K.M., Shimada, Y., Ayers, K., Lu, F., Hudson, A.M., and Cooley, L. (2015). Somatic insulin signaling regulates a germline starvation response in Drosophila egg chambers. Dev. Biol. 398, 206–217. 10.1016/j.ydbio.2014.11.021.

166. LaFever, L., and Drummond-Barbosa, D. (2005). Direct control of germline stem cell division and cyst growth by neural insulin in Drosophila. Science 309, 1071–1073. 10.1126/science.1111410.

167. Richard, D.S., Rybczynski, R., Wilson, T.G., Wang, Y., Wayne, M.L., Zhou, Y., Partridge, L., and Harshman, L.G. (2005). Insulin signaling is necessary for vitellogenesis in Drosophila melanogaster independent of the roles of juvenile hormone and ecdysteroids: female sterility of the chico1 insulin signaling mutation is autonomous to the ovary. J. Insect Physiol. 51, 455–464. 10.1016/j.jinsphys.2004.12.013.

168. Chandegra, B., Tang, J.L.Y., Chi, H., and Alic, N. (2017). Sexually dimorphic effects of dietary sugar on lifespan, feeding and starvation resistance in Drosophila. Aging 9, 2521–2528. 10.18632/aging.101335.

169. Eichmann, T.O., and Lass, A. (2015). DAG tales: the multiple faces of diacylglycerol—stereochemistry, metabolism, and signaling. Cell. Mol. Life Sci. CMLS 72, 3931–3952. 10.1007/s00018-015-1982-3.

170. Antony, C., and Jallon, J.-M. (1982). The chemical basis for sex recognition in *Drosophila melanogaster*. J. Insect Physiol. 28, 873–880. 10.1016/0022-1910(82)90101-9.

171. Jallon, J.M. (1984). A few chemical words exchanged by Drosophila during courtship and mating. Behav. Genet. 14, 441–478. 10.1007/BF01065444.

172. Fedina, T.Y., Kuo, T.-H., Dreisewerd, K., Dierick, H.A., Yew, J.Y., and Pletcher, S.D. (2012). Dietary Effects on Cuticular Hydrocarbons and Sexual Attractiveness in Drosophila. PLoS ONE 7, e49799. 10.1371/journal.pone.0049799.

173. Wicker-Thomas, C., Garrido, D., Bontonou, G., Napal, L., Mazuras, N., Denis, B., Rubin, T., Parvy, J.-P., and Montagne, J. (2015). Flexible origin of hydrocarbon/pheromone precursors in Drosophila melanogaster. J. Lipid Res. 56, 2094– 2101. 10.1194/jlr.M060368.

174. Scheitz, C.J.F., Guo, Y., Early, A.M., Harshman, L.G., and Clark, A.G. (2013). Heritability and Inter-Population Differences in Lipid Profiles of Drosophila melanogaster. PLOS ONE 8, e72726. 10.1371/journal.pone.0072726.

175. Faust, J.E., Manisundaram, A., Ivanova, P.T., Milne, S.B., Summerville, J.B., Brown, H.A., Wangler, M., Stern, M., and McNew, J.A. (2014). Peroxisomes Are Required for Lipid Metabolism and Muscle Function in Drosophila melanogaster. PLOS ONE 9, e100213. 10.1371/journal.pone.0100213.

176. Lodhi, I.J., and Semenkovich, C.F. (2014). Peroxisomes: a Nexus for Lipid Metabolism and Cellular Signaling. Cell Metab. 19, 380–392. 10.1016/j.cmet.2014.01.002.

177. Pridie, C., Ueda, K., and Simmonds, A.J. (2020). Rosy Beginnings: Studying Peroxisomes in Drosophila. Front. Cell Dev. Biol. 8. 10.3389/fcell.2020.00835.

178. Tuthill, B.F., Searcy, L.A., Yost, R.A., and Musselman, L.P. (2020). Tissue-specific analysis of lipid species in Drosophila during overnutrition by UHPLC-MS/MS and MALDI-MSI [S]. J. Lipid Res. 61, 275–290. 10.1194/jlr.RA119000198.

179. Santoro, C., O’Toole, A., Finsel, P., Alvi, A., and Musselman, L.P. (2022). Reducing ether lipids improves Drosophila overnutrition-associated pathophysiology phenotypes via a switch from lipid storage to beta-oxidation. Sci. Rep. 12, 13021. 10.1038/s41598-022-16870-4.

180. Alfa, R.W., Park, S., Skelly, K.-R., Poffenberger, G., Jain, N., Gu, X., Kockel, L., Wang, J., Liu, Y., Powers, A.C., et al. (2015). Suppression of Insulin Production and Secretion by a Decretin Hormone. Cell Metab. 21, 323–333. 10.1016/j.cmet.2015.01.006.

181. Luo, J., Lushchak, O.V., Goergen, P., Williams, M.J., and Nässel, D.R. (2014). Drosophila Insulin-Producing Cells Are Differentially Modulated by Serotonin and Octopamine Receptors and Affect Social Behavior. PLOS ONE 9, e99732. 10.1371/journal.pone.0099732.

182. Söderberg, J.A.E., Carlsson, M.A., and Nässel, D.R. (2012). Insulin-Producing Cells in the Drosophila Brain also Express Satiety-Inducing Cholecystokinin-Like Peptide, Drosulfakinin. Front. Endocrinol. 3, 109. 10.3389/fendo.2012.00109.

183. Yurgel, M.E., Kakad, P., Zandawala, M., Nässel, D.R., Godenschwege, T.A., and Keene, A.C. (2019). A single pair of leucokinin neurons are modulated by feeding state and regulate sleep–metabolism interactions. PLoS Biol. 17, e2006409. 10.1371/journal.pbio.2006409.

184. Kwak, S.-J., Hong, S.-H., Bajracharya, R., Yang, S.-Y., Lee, K.-S., and Yu, K. (2013). Drosophila Adiponectin Receptor in Insulin Producing Cells Regulates Glucose and Lipid Metabolism by Controlling Insulin Secretion. PLOS ONE 8, e68641. 10.1371/journal.pone.0068641.

185. Buch, S., Melcher, C., Bauer, M., Katzenberger, J., and Pankratz, M.J. (2008). Opposing Effects of Dietary Protein and Sugar Regulate a Transcriptional Target of Drosophila Insulin-like Peptide Signaling. Cell Metab. 7, 321–332. 10.1016/j.cmet.2008.02.012.

186. Lee, G., Bahn, J.H., and Park, J.H. (2006). Sex- and clock-controlled expression of the neuropeptide F gene in Drosophila. Proc. Natl. Acad. Sci. 103, 12580–12585. 10.1073/pnas.0601171103.

187. Asahina, K., Watanabe, K., Duistermars, B.J., Hoopfer, E., González, C.R., Eyjólfsdóttir, E.A., Perona, P., and Anderson, D.J. (2014). Tachykinin-expressing neurons control male-specific aggressive arousal in Drosophila. Cell 156, 221–235. 10.1016/j.cell.2013.11.045.

188. Shankar, S., Chua, J.Y., Tan, K.J., Calvert, M.E.K., Weng, R., Ng, W.C., Mori, K., and Yew, J.Y. (2015). The neuropeptide tachykinin is essential for pheromone detection in a gustatory neural circuit. eLife 4, e06914. 10.7554/eLife.06914.

189. Kim, W.J., Jan, L.Y., and Jan, Y.N. (2013). A PDF/NPF neuropeptide signaling circuitry of male Drosophila melanogaster controls rival-induced prolonged mating. Neuron 80, 1190–1205. 10.1016/j.neuron.2013.09.034.

190. Liu, C., Zhang, B., Zhang, L., Yang, T., Zhang, Z., Gao, Z., and Zhang, W. (2020). A neural circuit encoding mating states tunes defensive behavior in Drosophila. Nat. Commun. 11, 3962. 10.1038/s41467-020-17771-8.

191. Lebreton, S., Mansourian, S., Bigarreau, J., and Dekker, T. (2016). The Adipokinetic Hormone Receptor Modulates Sexual Behavior, Pheromone Perception and Pheromone Production in a Sex-Specific and Starvation-Dependent Manner in Drosophila melanogaster. Front. Ecol. Evol. 3. 10.3389/fevo.2015.00151.

192. Rezával, C., Nojima, T., Neville, M.C., Lin, A.C., and Goodwin, S.F. (2014). Sexually dimorphic octopaminergic neurons modulate female postmating behaviors in Drosophila. Curr. Biol. CB 24, 725–730. 10.1016/j.cub.2013.12.051.

193. Wang, T., Jing, B., Deng, B., Shi, K., Li, J., Ma, B., Wu, F., and Zhou, C. (2022). Drosulfakinin signaling modulates female sexual receptivity in Drosophila. eLife 11, e76025. 10.7554/eLife.76025.

194. Ameku, T., Yoshinari, Y., Texada, M.J., Kondo, S., Amezawa, K., Yoshizaki, G., Shimada-Niwa, Y., and Niwa, R. (2018). Midgut-derived neuropeptide F controls germline stem cell proliferation in a mating-dependent manner. PLOS Biol. 16, e2005004. 10.1371/journal.pbio.2005004.

195. Zhang, C., Daubnerova, I., Jang, Y.-H., Kondo, S., Žitňan, D., and Kim, Y.-J. (2021). The neuropeptide allatostatin C from clock-associated DN1p neurons generates the circadian rhythm for oogenesis. Proc. Natl. Acad. Sci. U. S. A. 118, e2016878118. 10.1073/pnas.2016878118.

196. Lin, W.-S., Yeh, S.-R., Fan, S.-Z., Chen, L.-Y., Yen, J.-H., Fu, T.-F., Wu, M.-S., and Wang, P.-Y. (2018). Insulin signaling in female Drosophila links diet and sexual attractiveness. FASEB J. 32, 3870–3877. 10.1096/fsb2fj201800067r.

197. Cong, X., Wang, H., Liu, Z., He, C., An, C., and Zhao, Z. (2015). Regulation of Sleep by Insulin-like Peptide System in Drosophila melanogaster. Sleep 38, 1075–1083. 10.5665/sleep.4816.

198. Isabel, G., Martin, J.-R., Chidami, S., Veenstra, J.A., and Rosay, P. (2005). AKH-producing neuroendocrine cell ablation decreases trehalose and induces behavioral changes in Drosophila. Am. J. Physiol. Regul. Integr. Comp. Physiol. 288, R531–538. 10.1152/ajpregu.00158.2004.

199. Bednářová, A., Tomčala, A., Mochanová, M., Kodrík, D., and Krishnan, N. (2018). Disruption of Adipokinetic Hormone Mediated Energy Homeostasis Has Subtle Effects on Physiology, Behavior and Lipid Status During Aging in Drosophila. Front. Physiol. 9, 949. 10.3389/fphys.2018.00949.

200. Post, S., Liao, S., Yamamoto, R., Veenstra, J.A., Nässel, D.R., and Tatar, M. (2019). Drosophila insulin-like peptide dilp1 increases lifespan and glucagon-like Akh expression epistatic to dilp2. Aging Cell 18, e12863. 10.1111/acel.12863.

201. Stephenson, E.J., Stayton, A.S., Sethuraman, A., Rao, P.K., Meyer, A., Gomes, C.K., Mulcahy, M.C., McAllan, L., Puchowicz, M.A., Pierre, J.F., et al. (2022). Chronic intake of high dietary sucrose induces sexually dimorphic metabolic adaptations in mouse liver and adipose tissue. Nat. Commun. 13, 6062. 10.1038/s41467-022-33840-6.

202. Mauvais-Jarvis, F., Merz, N.B., Barnes, P.J., Brinton, R.D., Carrero, J.-J., DeMeo, D.L., Vries, G.J.D., Epperson, C.N., Govindan, R., Klein, S.L., et al. (2020). Sex and gender: modifiers of health, disease, and medicine. The Lancet 396, 565–582. 10.1016/S0140-6736(20)31561-0.

203. Mauvais-Jarvis, F. (2015). Sex differences in metabolic homeostasis, diabetes, and obesity. Biol. Sex Differ. 6, 14. 10.1186/s13293-015-0033-y.

204. Chella Krishnan, K., Mehrabian, M., and Lusis, A.J. (2018). Sex differences in metabolism and cardiometabolic disorders. Curr. Opin. Lipidol. 29, 404–410. 10.1097/MOL.0000000000000536.

205. Tramunt, B., Smati, S., Grandgeorge, N., Lenfant, F., Arnal, J.-F., Montagner, A., and Gourdy, P. (2020). Sex differences in metabolic regulation and diabetes susceptibility. Diabetologia 63, 453–461. 10.1007/s00125-019-05040-3.

206. Strack, C., Behrens, G., Sag, S., Mohr, M., Zeller, J., Lahmann, C., Hubauer, U., Loew, T., Maier, L., Fischer, M., et al. (2022). Gender differences in cardiometabolic health and disease in a cross-sectional observational obesity study. Biol. Sex Differ. 13, 8. 10.1186/s13293-022-00416-4.

207. Veenstra, J.A., Agricola, H.-J., and Sellami, A. (2008). Regulatory peptides in fruit fly midgut. Cell Tissue Res. 334, 499–516. 10.1007/s00441-008-0708-3.

208. Kasturacharya, N., Dhall, J.K., and Hasan, G. (2023). A STIM dependent dopamine-neuropeptide axis maintains the larval drive to feed and grow in Drosophila. PLOS Genet. 19, e1010435. 10.1371/journal.pgen.1010435.

209. Schindelin, J., Arganda-Carreras, I., Frise, E., Kaynig, V., Longair, M., Pietzsch, T., Preibisch, S., Rueden, C., Saalfeld, S., Schmid, B., et al. (2012). Fiji: an open-source platform for biological-image analysis. Nat. Methods 9, 676–682. 10.1038/nmeth.2019.

210. Yu, H., and Huan, T. (2022). MAFFIN: metabolomics sample normalization using maximal density fold change with high-quality metabolic features and corrected signal intensities. Bioinformatics 38, 3429–3437. 10.1093/bioinformatics/btac355.

211. May, C.E., Vaziri, A., Lin, Y.Q., Grushko, O., Khabiri, M., Wang, Q.-P., Holme, K.J., Pletcher, S.D., Freddolino, P.L., Neely, G.G., et al. (2019). High Dietary Sugar Reshapes Sweet Taste to Promote Feeding Behavior in Drosophila melanogaster. Cell Rep. 27, 1675–1685.e7. 10.1016/j.celrep.2019.04.027.

212. Nunes, R.D., and Drummond-Barbosa, D. (2023). A high-sugar diet, but not obesity, reduces female fertility in Drosophila melanogaster. Dev. Camb. Engl. 150, dev201769. 10.1242/dev.201769.

213. Li, Y., Wang, W., and Lim, H.-Y. (2021). Drosophila Solute Carrier 5A5 Regulates Systemic Glucose Homeostasis by Mediating Glucose Absorption in the Midgut. Int. J. Mol. Sci. 22, 12424. 10.3390/ijms222212424.zf

214. LeDue, E.E., Mann, K., Koch, E., Chu, B., Dakin, R., and Gordon, M.D. (2016). Starvation-Induced Depotentiation of Bitter Taste in Drosophila. Curr. Biol. CB 26, 2854– 2861. 10.1016/j.cub.2016.08.028.

215. Marella, S., Mann, K., and Scott, K. (2012). Dopaminergic modulation of sucrose acceptance behavior in Drosophila. Neuron 73, 941–950. 10.1016/j.neuron.2011.12.032.

216. Wobbrock, J. O., Findlater, L., Gergle, D., & Higgins, J. J. (2011). The aligned rank transform for nonparametric factorial analyses using only anova procedures. Proceedings of the SIGCHI Conference on Human Factors in Computing Systems, 143–146. 10.1145/1978942.1978963

