## Supplementary figures and images for "Insulin/insulin-like growth factor signaling pathway promotes higher fat storage in *Drosophila* females"

### Supplemental Figures

#### Supplemental Figure 1

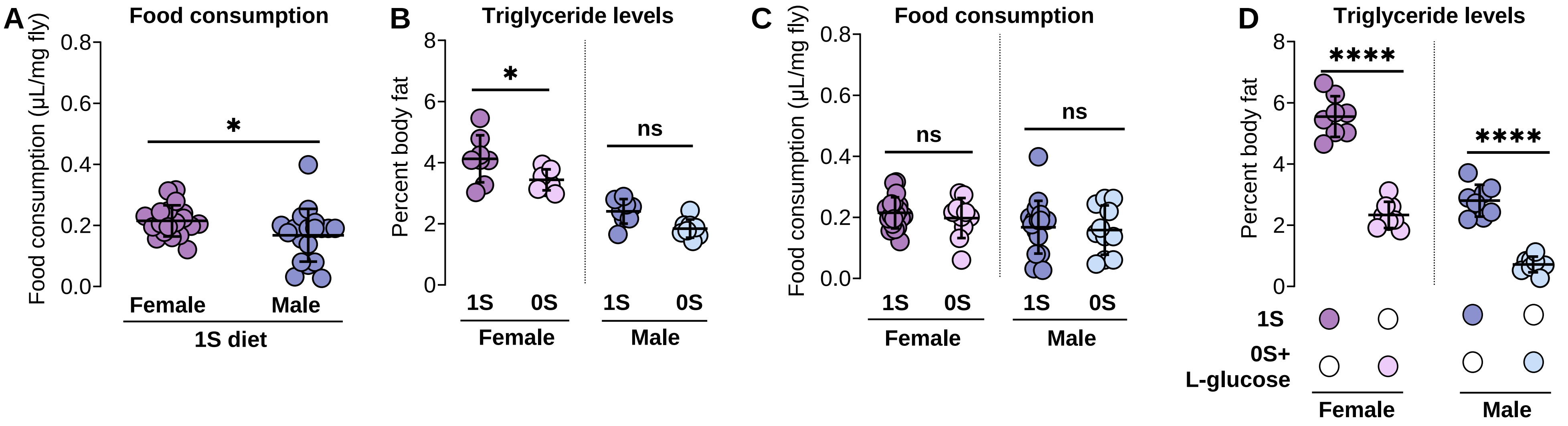

##### Supplemental Figure 2

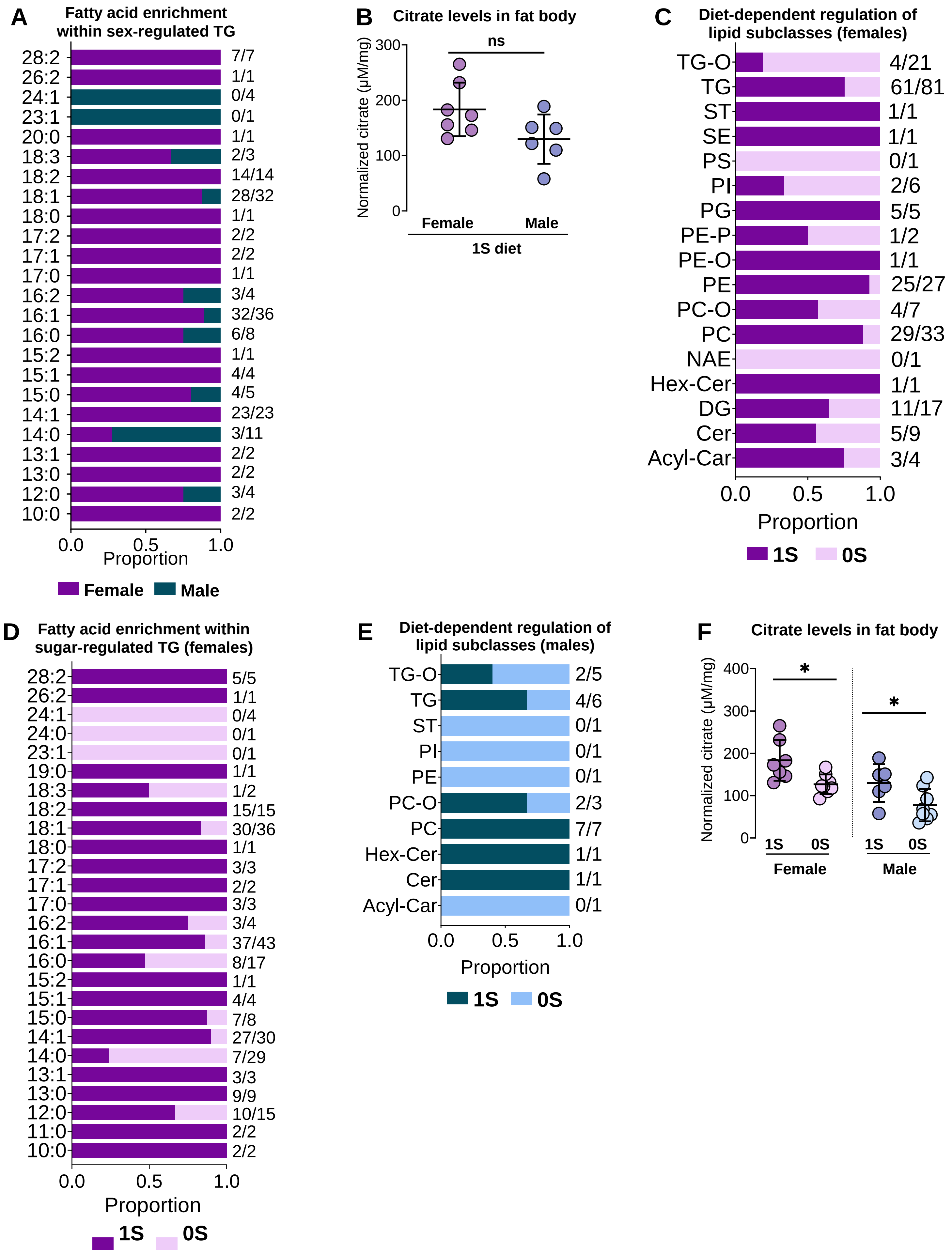

#### Supplemental Figure 3

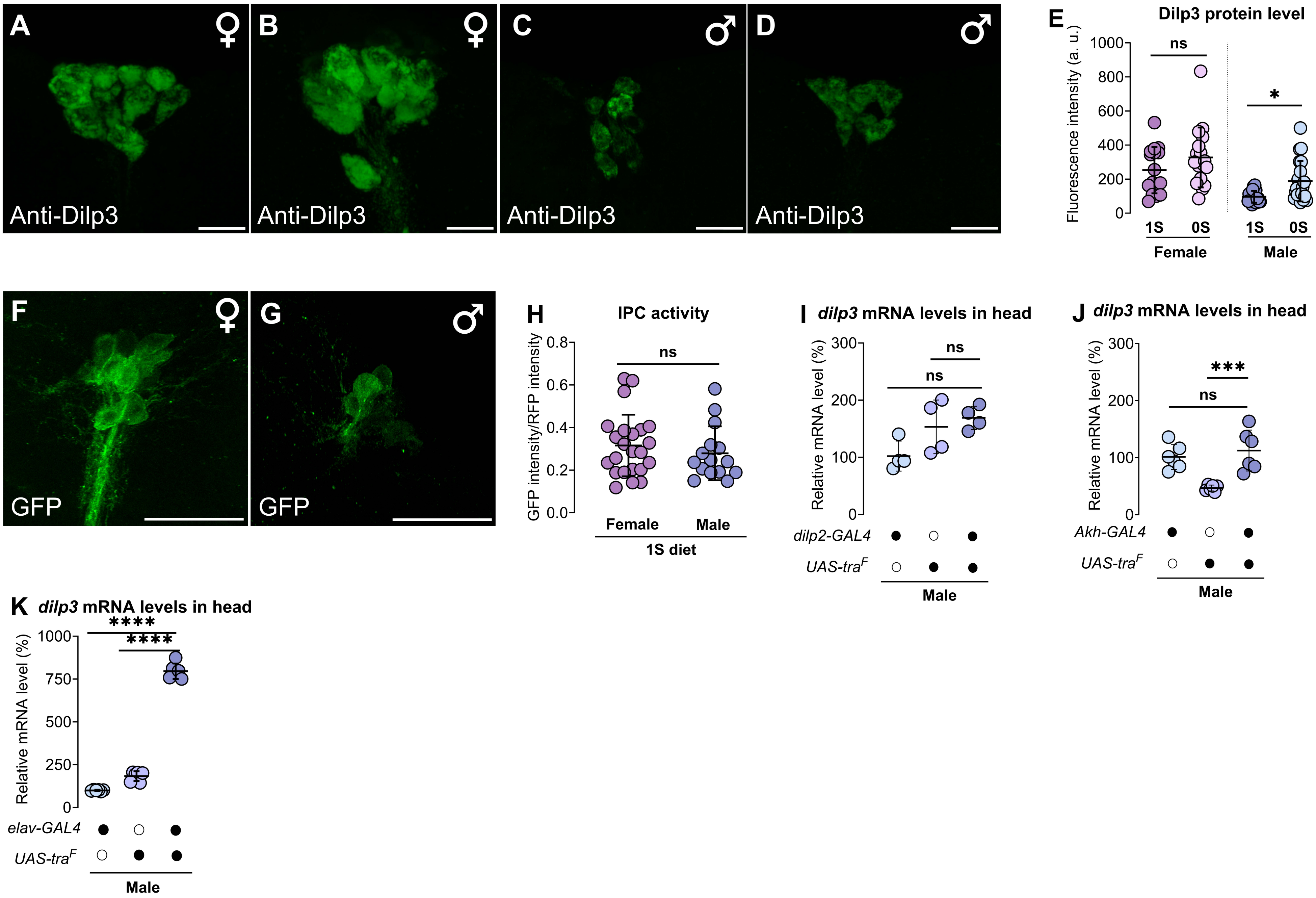

#### Supplemental Figure 4

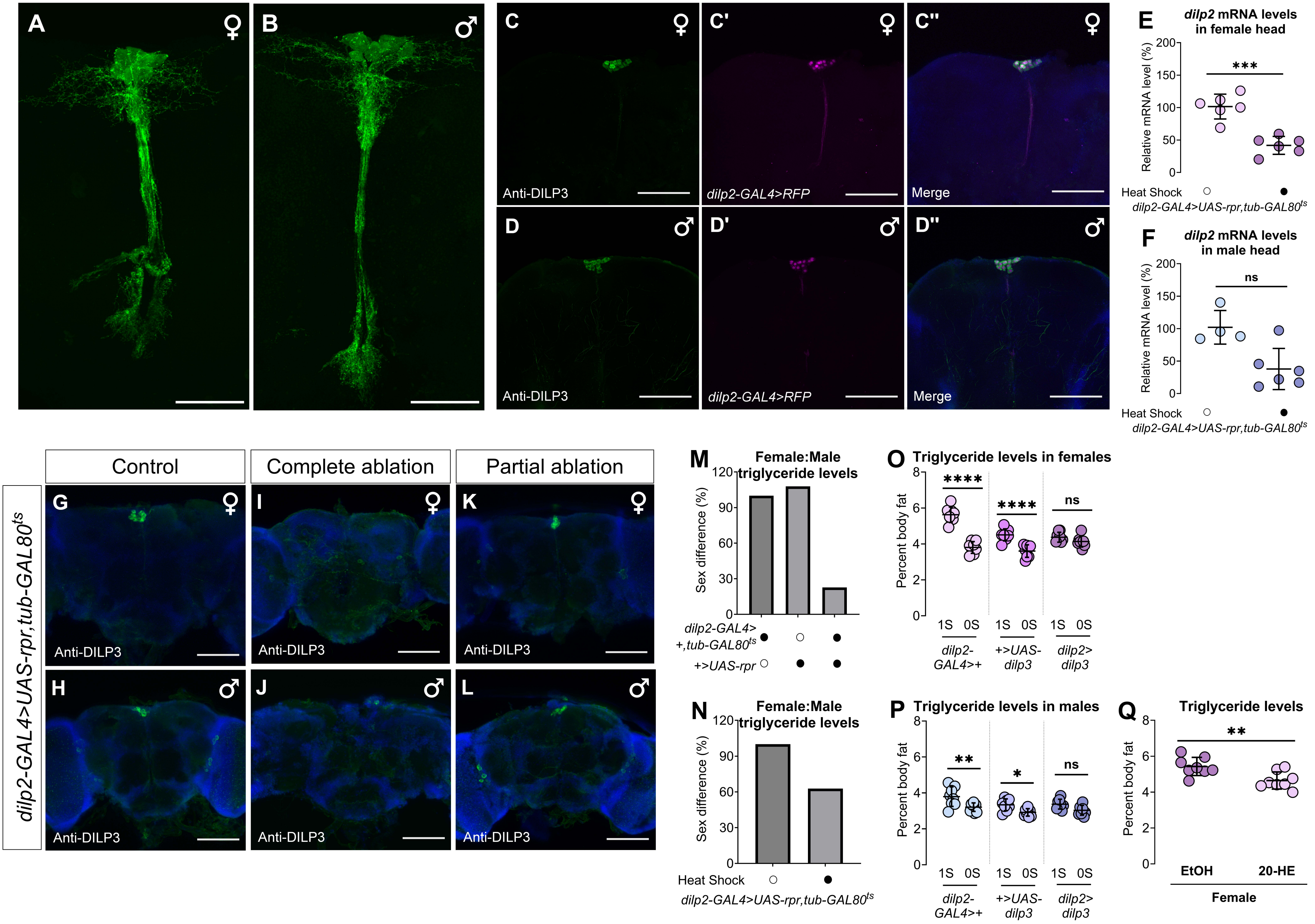
